# Microbial Response to Natural Disturbances: Rare Biosphere often plays a role

**DOI:** 10.1101/2024.03.06.583742

**Authors:** Jianshu Zhao, Genevieve Brandt, Zhao Wang, Dana E. Hunt, Luis M. Rodriguez-R, Janet K. Hatt, Konstantinos T. Konstantinidis

## Abstract

Understanding how microbial populations respond to disturbances represents a major goal for microbial ecology. While several theories have been advanced to explain microbial community compositional changes in response to disturbances, appropriate data to test these theories is scarce, especially when considering the challenges to define rare vs. abundant taxa and generalists vs. specialists, a prerequisite for testing the theories. Here, we define these two key concepts by employing the patterns of coverage of a (target) genome by a metagenome to define rare populations, and by borrowing concepts from macroecology, the proportional similarity index (PS index), to define generalists. Using these concepts, we found that coastal microbial communities are resilient to major perturbations such as tropical cyclones and (uncommon) cold or warm weather events snaps-in part-due to the response of rare populations, providing support for the insurance hypothesis (i.e., the rare biosphere has the buffering capacity to mitigate the effects of disturbances). Generalists appear to contribute proportionally more than specialists to community adaptation to perturbations like warming, supporting the disturbance-specialization hypothesis, i.e., disturbance favors generalists. Taken together, our results advance understanding of the mechanisms governing microbial populations dynamics under changing environmental conditions and have potential applications for ecosystem management.

## Introduction

Natural environmental microbial communities generally harbor a few abundant taxa and numerous low abundance, or rare, taxa [1]. In recent years, the importance of the rare biosphere has been increasingly emphasized. Rare biosphere is defined as the low-abundance active or dormant microbial taxa in a given environment at a specific time point, typically showing <0.1% relative abundance. Low-abundance taxa are important contributors to both α- and β-diversity, and rare taxa have been often shown to have important ecological roles in specific ecosystems [2]. Both the number of taxa (phylogenetic diversity) and the genes these taxa carry (functional diversity) are thought to provide an ‘insurance’ or ‘buffering/caching’ capacity for the ecosystem against environmental change [3,4]. For example, a rare coastal, oil-degrading bacterial population thrived (e.g., made up ∼30% of the total community) three weeks after an oil spill and became rare two months later, after the oil was degraded [5]. Members of the rare biosphere have also been shown to become abundant in a bacterioplankton community after disturbance and play important roles in maintaining ecosystem processes [6]. Rare taxa can periodically become abundant, and their abundance dynamics may depend on selection pressure such as those imposed by seasonal fluctuations in environmental parameters and substrate availability, and/or ecological processes including dispersal, predation, reactivation from dormancy, functional redundancy, plasticity, and diversification [7,8].

Several ecological theories have been advanced to explain the responses of abundant and rare microbial populations to disturbance. For example, the insurance hypothesis suggests that rare biosphere represents a strategy for responding to temporally variable environments, contributing to community resilience [9,10]. Specifically, the hypothesis predicts that rare species, probably beyond the limit of detection and thus, not included in estimates of community diversity, may quickly respond to altered environmental characteristics (pulse disturbance) and become abundant before returning to pre-perturbation (low) abundances [11]. On the other hand, the specialization-disturbance hypothesis suggests that niche breadth, rather than relative abundance, of a species is important for how the species responds to disturbance. That is, disturbance is usually expected to affect specialists negatively, while generalists are thought to benefit from disturbance [12]. Defining generalist versus specialist taxa in environmental samples relies on their prevalence/abundance or niche width: generalists generally show no preference for specific environments contrasting with specialists, which are abundant only in some environments or conditions. Both theoretical and laboratory research has shown that generalist taxa can be crucial in maintaining ecosystem functioning under fluctuating/disturbed environments compared to specialists due to their metabolic flexibility [13,14], although defining generalist vs. specialists is practically challenging due to difficulties in defining the niche breadth that each species may occupy and determining relative abundance when this is low (see also below).

Although the rare biosphere often plays an essential role in community function and stability, defining rare and abundant members has largely employed arbitrary divisions. Several different relative abundance thresholds have been proposed (e.g., <1%, 0.1% or 0.01% of the total community), without attention to ecological theory or habitat differences [15–17]. Moreover, according to random sampling theory, defining rare microbial taxa in comparison to abundant taxa based on 16S rRNA gene amplicon sequence data can be subjected to overestimates of relative abundance due to amplification biases (e.g. abundant populations are favored during amplification and/or sequencing) [18]. More importantly, amplicon sequencing provides little information on population-level functional potential and thus, offers limited information on the roles of abundant and rare taxa in maintaining important ecosystem processes. Metagenomics provides the means to capture the functional diversity of rare biosphere and largely sidestep the biased amplification of abundant taxa based on gene-amplicon data [19,20] but it remains challenging to reconstruct the metagenome-assembled genomes (MAGs) of rare taxa due to insufficient sampling (i.e., sequence coverage), and to define rare vs. abundant taxa at the genome level [21].

In this study, we sought to quantify how often rare populations respond to disturbances and what functions rare populations provide relative to abundant taxa. In other words, we aimed to directly test the specialization-disturbance and the insurance hypotheses. For this, we sampled surface water at a costal observatory (PICO) at Beaufort, North Carolina (USA) weekly, for a three-year period, during which disturbance events occurred (initially identified by microbiome changes, see below), with at least 1 to 3 weeks of season-typical weather between these events (Figure S1). Disturbance were defined previously by comparing whether overall community composition and structure (e.g. beta diversity) changed by the event (e.g., cyclone, bloom) relative to the composition that is typical for the same season that the event occurred [22]. Shotgun metagenomes before, during and after these events were used to define abundant and rare population based on MAG coverage (e.g., genome depth and breadth covered by mapped reads) and assess how many rare populations responded to the events. In this process, we also defined the limit of detection of our metagenomic effort based on the species abundance distribution curve as well as generalists vs. specialists based on a proportional similarity index (PS index). Therefore, this study not only quantifies the importance of the rare biosphere during community response to disturbance but also outlines a methodology to define challenging concepts for microbial ecology, such as rare versus abundant population and specialists versus generalists.

## Materials and Methods

### Sample collection and description of disturbance events

Water samples were collected as part of the Piver’s Island Coastal Observatory (PICO) time series adjacent to the Beaufort Inlet, Beaufort, NC, USA (34°71.81’N, 76°67.08’W) on a weekly basis [22]. Here, we sequenced 19 samples from January 2011 to December 2013 that represented the winter, summer, and spring seasons. For each season in each year, 3 samples were sequenced representing non-disturbance, disturbance, and post-disturbance samples of disturbance events. Disturbance events were identified as significant changes compared to the seasonal microbial community composition and structure based on 16S rRNA gene amplicon sequencing [22] (**Figure S1**, Table S1). We also looked at the actually environmental parameters that might have driven those changes among the parameters measured. Briefly, for the six disturbance events identified previously by Gronniger *et al.* [22], disturbance 1 was a warm and windy week in winter of 2011 (50% less ammonium in the disturbance sample relative to non-disturbance samples), disturbance 2 was the cyclone in summer of 2011 (Hurricane Irene, 65% more ammonium, 80.2% more silicates and 78.2% more chlorophyll A), disturbance 3 was a rainy week in winter 2012 (10 times lower than the two low tides, 3 times more ammonium), disturbance 5 was a warm week in spring of 2012 (116% more NOx, 55.9% less silicates and 31.7% less chlorophyll A), disturbance 8 was a warm week in spring 2013 (100.1% more chlorophyll A and 59% less ammonium), disturbance 9 was the tropical cyclone in summer 2013 (220.3% more ammonium, 49.6% more silicates and 20.2% more chlorophyll A, 1.5 ^0^C higher). We refer to those disturbance events as win11_warm_1, sum11_cyclone_2, win12_rainy_3, spr12_warm_5, spr13_warm_8, sum13_cyclone_9, respectively, with the last number corresponding to the disturbance events label shown in Figure S1 and elsewhere, which is also consistent with our previous publication [22]. Seawater was collected at 10:30 h local time using a 5-liter Niskin bottle centered at 1 m on a peristaltic pump with the tubing open at 1 m and processed within 1 h. Standard laboratory methods for determination of water temperature, pH, salinity, dissolved inorganic nutrient concentrations, and chlorophyll-a concentrations were described previously [23]

### DNA extraction, amplicon sequencing analysis

Microbial biomass was collected by filtering ∼1 liter of seawater through a 0.22-micron Sterivex filter (Millipore, Darmstadt, Germany) and the filters were stored at −80 °C until DNA extraction. Genomic DNA was extracted using the phenol-chloroform lysis supplemented with bead beating (60 seconds) and then subsequently cleaned using the Zymo OneStep PCR inhibitor removal kit. Extracted DNA was quantified using a Nanodrop ND-100 before sequencing. 16S rRNA gene amplicons from each sample were sequenced using the primers targeting the V3-to-V4 region of the bacterial and archaeal 16S rRNA genes: for 16S F V3, CCTACGGGNGGCWSCAG; and for 16S R V4, GGACTACNVGGGTWTCTAAT as described previously [24]. PCR reactions contained 10 ng of template DNA, 1.25 U Econo Taq (Lucigen) and a final concentration of 1× Taq buffer, 200 μM dNTPs, 2 mM MgCl_2_ and 0.5 μM of each primer. PCR reactions were performed with the following protocol: 98°C for 30 seconds followed by 35 cycles at 98°C for 10 seconds, then 55°C for 30 seconds and 72°C for 30 seconds, with a final extension at 72°C for 2 minutes. Triplicate reaction mixtures per sample were pooled and gel purified. Paired-end 250-bp sequencing of barcoded amplicons was performed on an Illumina MiSeq running v2 chemistry at the Duke Center for Genomic and Computational Biology.

USEARCH v11.0.667 was used for quality control and merging of paired-end reads. We first trimmed low-quality bases from the sequences using a 10-nucleotide (nt) window with a Q30 running-quality threshold. Paired-end sequences with a >=10-nt overlap and no mismatches were then merged. We performed a final filtering step to discard low-quality merged sequences with a length of <400 bp and/or a maximum expected error of >1. Amplicon sequence variants (ASVs) were then identified using the UNOISE3 algorithm in USEARCH [25], which has been shown to be even more accurate than DADA2 [26]. Taxonomy classification of representative ASV sequences was performed using SINTAX (-sintax_cutoff 0.8, a significance threshold similar to the 50% bootstrap cutoff accuracy of the RDP naïve classifier) against SILVA v138.1 in USEARCH [27,28]. MacQIIME v1.9.1 was used for rarefaction, alpha diversity, beta diversity and community composition analysis [29].

### Metagenomic sequencing, quality control and coverage estimation

DNA was extracted with the MoBio Power Soil kit (MoBio Inc.Carlsbad, CA, USA). 10 ng of DNA was then sheared to 300 bp using the Covaris LE220 and size was selected using SPRI beads (Beckman Coulter). The fragments were treated with end-repair, A-tailing, and ligation of Illumina compatible adapters using the KAPA-Illumina library creation kit followed by 5 cycles of PCR to enrich for the final library. These libraries were sequenced with 2 150 nt reads on the Illumina HiSeq 2500 1T platform at either Duke’s Genome Sequencing or the Department of Energy’s Joint Genome Institute for 300 cycles. Adapter trimming and demultiplexing of sequenced samples was carried out by the instrument. Raw reads were trimmed using Trimmomatic with default parameters [30] and then checked using FastQC (https://github.com/s-andrews/FastQC). Sequence subsampling to account for sequencing depth variation among libraries was done using Seqtk and specifically, the subseq command with the same random seed for forward and reverse reads (https://github.com/lh3/seqtk). Sequencing coverage estimation and Nonpareil diversity were calculated using Nonpareil v3.0 with-kmer option [31].

### 16S rRNA gene-carrying read extraction from metagenomes and closed reference OTU picking

Prokaryotic 16S rRNA gene-carrying metagenomic reads were extracted using metaxa2 [32]. USEARCH closed_ref workflow in USEARCH v11.0.667 were performed to pick closed reference OTUs (not ASVs since we cannot cluster 16S short sequences extracted from metagenome) from the extracted 16S rRNA gene-carrying (16S) short reads with default identity 97% [29,33]. Briefly, USEARCH was used to align extracted short 16S rRNA gene reads against the non-redundant SILVA database v138.1. If the semi-global alignment identity of a query sequence to a database sequence was better than 97%, the input sequence was assigned to that OTU. Otherwise, the input sequence was discarded and not assigned to an OTU (an identity threshold of 99% was also used but only small differences were observed in the results in terms of community composition, especially at higher than the family taxonomic levels). Extracted 16S rRNA gene reads were rarefied to 8000 sequences/sample before downstream analysis (Figure S3). Note that closed reference OTU clustering may not find an OTU for a (query) sequence that is not represented with a closely related sequence in the SILVA database, contrasting with de novo OTU/ASV clustering using amplicons. However, such sequences were rather rare in our datasets, representing <5% of the total sequences for each sample; hence, no further action was taken to deal with this issue. Downstream analyses including diversity and compositions analysis were performed using MacQIIME v1.9.1.

### Metagenome assembly, contig taxonomic classification, coverage calculation and functional gene diversity analysis

Quality controlled short reads were assembled with Megahit v2.1.2 (parameters: --meta - -min_contig 1000) [34]. To classify the assembled contigs taxonomically, Centrifuge was used to search against the RefSeq complete genome collection with default parameters [35]. Genes were predicted on contigs using Prodigal v2.6.3 (-p meta) [36]. After mapping short reads onto each contig and the genes of a contig using bwa-mem2 [37], contig/gene coverage was calculated by the CoverM v0.6.1 (https://github.com/wwood/CoverM), contig workflow with abundance metric option metabat (--methods metabat --min-read-percent-identity 0.95 --min-read-aligned-percent 0.75). Diamond v0.9.22 was used to perform functional annotation of predicted genes against the Swiss-Prot database (Diamond blastp -k 1 --id 40 --query-cover 70 --max-hsps 4 -e 0.0001) [38]. The matching Swiss-Prot reference sequences were mapped to GO terms and filtered for molecular functions. The Chao-Shen estimate of Shannon entropy H for molecular functions and cellular processes was calculated using the R package Entropy based on the observed read counts (coverage) of genes assigned to the same molecular function GO term. This estimation adjusts for missing species (here GO terms) and sample coverage. The exponential of the estimated Shannon entropy was used to covert the statistic to true diversity (^1^D) with units of effective GO terms as described recently [39]. The fraction of the total proteome devoted to extracellular proteins for each MAG was predicted using psortb [40].

### Population genome binning, and bin/MAG refinement

Maxbin 2, Metabat2 and CONCOCT were individually applied to contig binning with default parameters [41–43]. Resulting MAGs were first refined (quality improvement) using DAS Tools with searching engine USEARCH [33,44]. Refined MAGs were further checked by CheckM unique command, to ensure that no contigs were binned to the same MAG more than once and one contig was not binned to different MAGs. Mis-binned contigs were removed from MAGs. Contig coverage depth and tetranucleotide frequency of each MAG were subsequently evaluated to determine whether binned MAGs likely represent chimeric sequences using an in-house R script (available at https://github.com/jianshu93/bin_check). CheckM lineage_wf was then used for MAG quality assessment with default parameters [45]. The quality score was defined as completeness - 5*contamination + Stain heterogeneity*0.5. Medium to high quality MAGs were defined as quality score larger than 0.5. To further check whether those MAGs binned from each sample represent sequence-discrete populations (species) [46], competitive read mapping on the MAGs followed by recruitment plot were performed via the scripts in the enveomics R package [47] (a new pipeline based on the scripts is also available here: https://github.com/jianshu93/RecruitmentPlot_blast). Briefly, contig sequences of all MAG from the same sample were labeled and then pooled together as one genome database. Subsequently, blastn search was performed to map quality-controlled short reads to the contigs (-task blastn -id 95% -max_target_seqs 500). The blastn tabular output was filtered to remove mapped reads with low alignment ratio (<90% of read length) and to only keep the best match for each query read according to identity.

Tied matches were also removed before creating recruitment plot using the BlastTab.recplots2.R. MiGA quality_wf with MyTaxa option was then performed to further validate the taxonomic identity of contigs binned into MAGs [48] and also calculate gene coding density, %G+C content, and other descriptive statistics such as contig length [49].

### MAG dereplication and classification

Dereplication of MAGs was performed using dRep v2.2.4 with an ANI threshold of 95% by the -fastANI option (minimum completeness 70%, maximum contamination 10% and quality score > 0.5 for filtering MAGs before dereplication) [50]. Quality information from CheckM was passed to dRep to select the best quality MAG as representative of each resulting 95% ΑΝΙ cluster. MAGs were classified against the Genome Taxonomy Database v214 [51] using GTDB-Tk classify_wf workflow [52], which classifies MAGs based on their placement in a reference tree inferred using pplacer analysis of a set of 120 bacterial and 122 archaeal concatenated gene markers, combined with FastANI for the species-level assignments [53,54]. To further confirm the taxonomy classified by GTDB-Tk, especially those that were not classified with confidence by GTDB-Tk (e.g., distantly related to their best matches found in GTDB), we also used the MiGA workflow miga classify_wf against the type material database [49]. The lowest taxonomic level to which MiGA considered the assignment of the query MAG as significant was kept and compared to the GTDB-Tk classification results.

### Relative abundance calculation and functional gene annotation of MAGs

The relative abundance of each dereplicated MAG was calculated by competitively mapping reads from each sample to the entire dereplicated MAG collection using bowtie2 and then SAMTools to generate sorted BAM files for each MAG [55,56]. Bam files were subsequently filtered using the CoverM workflow (--min-read-percent-identity 0.95 --min-read-aligned-percent 0.75) and only reads with alignment ratio and identity larger than 75% and 95%, respectively, were kept. Truncated Mean Depth 80% (TAD80) was then calculated as a proxy for relative DNA abundance, which normalizes for highly conserved or variable regions of the genome [57]. TAD80 estimates were further normalized by genome equivalents based on MicrobeCensus [58] to account for average genome size differences among the sample and provide the final (normalized) relative abundance estimates, using an in-house script (https://github.com/jianshu93/Competitive_mapping) [57]. Functional annotation of the dereplicated MAGs were performed using MicrobeAnnotator and DRAM [59,60]. Briefly, MicrobeAnnotator searches multiple reference protein databases iteratively and combines results from KEGG Orthology (KO), Enzyme Commission (E.C.), Gene Ontology (GO), Pfam and InterPro, and returns the matching annotations together with key metadata. DRAM profiles microbial (meta)genomes for metabolisms known to impact ecosystem function across biomes like carbon degradation, photosynthesis, methanogenesis, *etc*.

### Proportional similarity index for defining generalist and specialist MAGs

Levin’s niche breadth index (average relative abundance of species in different environments) has been recently used to define generalists vs. specialists for microbial populations [57,61,62]. However, this metric assumes equal availability of resources, which is hardly the case in natural environments. A metric based on the proportion similarity index, which defines generalists and specialists independently of their absolute abundance and occupancy, was used here to circumvent this limitation according to previous studies [63,64]. Specifically, habitat breadth was defined as the proportional similarity (PS) index [63] relating the proportion of a population (or species) found in each category of samples to the proportion of sampling effort (total samples) for that category as follows: 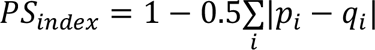, where p_i_ is the number of cells (we use relative abundance, i.e., TAD/genome equivalent) of the target species in samples of category i, divided by the total number of cells from that species in all available samples, while q_i_ is the number of all the cells in samples of category i, divided by all the cells in all samples. Thus, PS index values express variation in the habitat breadth of a species, which is an important aspect of the realized niche of the species (preferences for samples/environments), and range between 0 and 1 for the broadest possible and the narrowest possible niche, respectively (i.e., a population is restricted exclusively to the rarest category of sample types and consequently, is absent in all other types). Genomospecies captured in only one water sample were classified as rare species and excluded from the habitat breadth analysis as no reliable measure of niche breadth could be calculated for such genomospecies. For our time series dataset, samples were first grouped into three main types (or environments) according to seasonality, and then to two or three sub-types within each main type according to the year that the corresponding samples was obtained (Table S3). The PS index was calculated separately for each category of samples (the three main types and the eight sub-types), generating two scores (PS index_subtype_ & PS index_maintype_). Species that scored for both categories in the upper third percentile, indicating a broad niche, were classified as habitat generalists. Species that scored for both categories in the bottom third percentile were classified as habitat specialists, and were further subdivided into regular habitat specialists (rank-transformed; that is, the rank of the PS values, PS index_subtype_ > PS index_maintype_) and strict habitat specialists (rank-transformed; PS index_subtype_ < PS index_maintype_) [64](Table S3). According to this definition, species were classified as generalists or specialists independent of their relative abundance in one category of samples. This definition allows abundant (in some samples but not all) and widespread species (not equally abundant for all samples) to be also classified as habitat specialists when high proportions of individuals are found in a single category of samples (e.g., single season) but not in all category of samples [64]. We also compared this PS index with the Levin’s Breadth Index, and also the PS index implementation in MicroNiche package, which provides null model tests with respect to the limit of detection [65]. Specifically, for PS index by the MicroNiche, we used the principal component 1 of all available resource variables (e.g., DIC, Chlorophyll a, NH ^+^, NOx, PO ^3-^, SiO ^4-^) as the resource parameter in MicroNiche.

### Diversity analysis and statistics

All diversity and statistical analysis were performed using the R software (version 4.0.5) and its vegan, stats and ggplot2 packages for figures.

## Results

### Change in overall microbial community composition in response to disturbance events

Shifts in microbial community diversity did not show a consistent pattern between disturbance events. The Chao1 index based on either extracted 16S rRNA gene (or 16S) fragments recovered in metagenomes or 16S amplicon sequences neither increased nor decreased systematically by the winter disturbance events (win11_warmer_1, win12_windier_3) (Figure 1 (a)). Summer disturbance events (sum11_cyclone_2, sum13_cyclone_9), which represented cyclone events in year 2011 and 2013, either increased or decreased diversity despite representing similar weather changes (Figure 1). Nonpareil diversity, a reference database-free metric combining richness and evenness based on shotgun metagenomic data, also showed similar patterns to the Chao1 results reported above (Figure 1(b)). A consistent pattern was observed that, no matter whether disturbances increased or decreased diversity, diversity recovered after each disturbance, indicating that the sampled microbial communities were resilient. The resilience was also supported by community structure analysis based on NMDS of Mash distances of whole metagenomes, or amplicon and extracted metagenomic reads carrying 16S fragments (Figure 1(c) and Figure S4). Functional diversity analysis showed that molecular function diversity showed limited change with disturbance despite substantial diversity and composition changes (Figure S6), indicating high functional redundancy among the sampled microbial communities.

**Figure 1.**
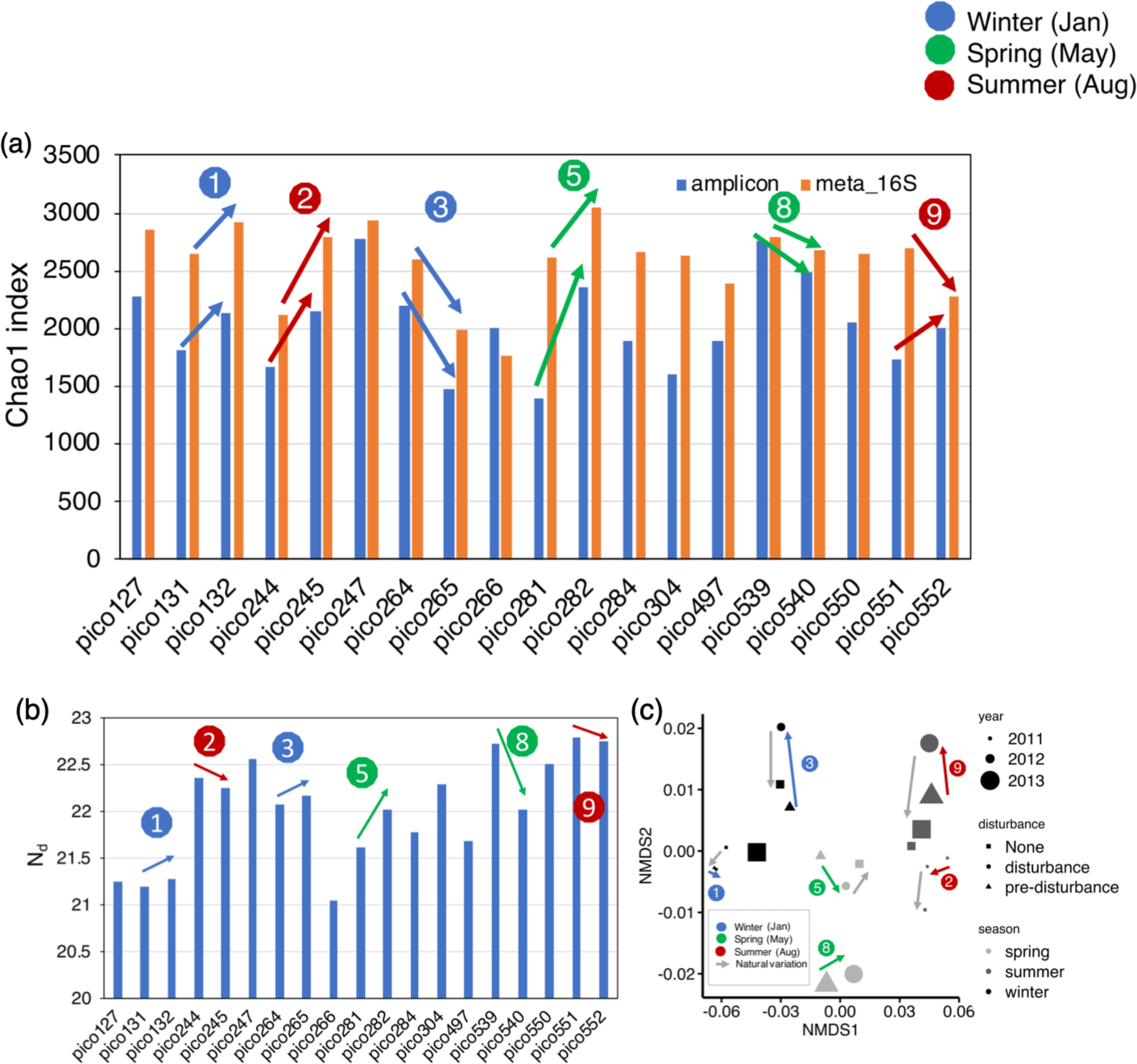
(a) Diversity shifts during disturbance events. Chao1 diversity index (a) for both amplicon 16S rRNA genes (blue) and extracted 16S rRNA gene reads from metagenomes (orange) are shown. Nonpareil diversity (N_d_) (b) and Mash distance based NMDS (c) of subsampled metagenomes are also included. Disturbance events are labelled with number (**see Figure S1 for details**) and the color indicates seasonality. Colored arrows in (a), (b) and (c) showed how N_d_ diversity and metagenomic composition was changed by each disturbance event. Grey arrows in (c) shows the natural variation of metagenomic composition (recovery or natural variation).

### Identifying abundant vs rare microbial populations

A total of 394 medium or high-quality MAGs (i.e., quality score >0.5) were obtained after quality control and before de-replication at 95% ANI. For each sample, contigs of different MAG fell into separate clusters based on dimension reduction of contig information (i.e., contig coverage and kmer profile) using the mmgenome2 software (Figure S14), confirming that most MAGs were likely not chimeric and represent species clusters. Recruitment plot showed that each MAG represented a sequence discrete population within the sample that it was recovered from, e.g., reads mapping between 85%-95% nucleotide identity were rare compared with reads showing >95% identity to the MAG (Figure S15, left and right bottom panels). These results were consistent with previous findings suggesting recognizable prokaryotic species may exist for natural communities [46]. There were 198 non-redundant MAGs after dereplication. Among these, three archaea MAGs were found, belonging to the newly proposed family *Candidatus* Poseidonaceae (formerly subgroup MGIIa) and one MAG belonging to *Ca.* Thalassarchaeaceae (formerly subgroup MGIIb). Among the bacterial populations, although order *Pelagibacterales* was very abundant at the read level (9.6% to 22.5% of the total metagenome/community), only one *Pelagibacterales* MAG was recovered with medium quality (quality score 0.57). All other *Pelagibacterales* MAGs were low in completeness and were removed at the MAG quality control check step. However, at the contig level, 4.4% to 5.2% contigs were classified as *Ca.* Pelagibacter, consistent with high relative abundance of this group at the read level. Inability of current genome binning algorithms to handle high intra-population diversity and recover representative genomes has been recently noted for this microbial group [66]. Most other MAGs were assigned to the classes of *Flavobacteriia* (phylum *Bacteroidetes*; recently renamed to *Bacteroidota* (*21*), *Alphaproteobacteria* and *Gammaproteobacteria* (phylum *Proteobacteria/Pseudomonadota (33 and 17*) with a few MAGs assigned to classes of other phyla (Table S2). These dereplicated MAGs represented collectively 5.1% to 18.2% of the total metagenomic reads for different samples, consistent with the high overall diversity of our samples that renders assembly and binning processes limiting.

Next, we pooled the recovered 198 non-redundant MAGs and competitively mapped reads from each of the 19 samples to calculate MAG relative abundance in order to subsequently draw the (genomo-)species abundance distribution (i.e., the abundance of each MAG is plotted against its rank among all MAGs according to their relative abundance; Figure 2). We used the resulting species-abundance curve to examine if there is any natural discontinuity or inflection point that could be used to define abundant vs. rare taxa more reliably than previous arbitrary threshold in abundance (discussed above). The species abundance curve was modeled robustly by a log scale model (Figure 2, Figure S8 (a) and (b), R^2^=0.732∼0.902, *P* < 0.01), and we observed no area of discontinuity in the data. A sharp decrease in terms of curve slope was observed around 0.1% relative abundance, however (Figure S7). We also observed a corresponding sharp decrease in MAG breadth coverage (how much of the genome is covered by reads) for relative abundance around 0.1% (Figure 2 and S8 (a) and (b)). Specifically, breadth coverage was 90% or less at this level of relative abundance and decreased quickly for less abundant genomospecies. Based on these results, we defined MAGs as rare when they showed relative abundance less than 0.1% and a coverage breadth less than 90% in that sample; we adjusted this threshold on a per-sample basis, depending on the sample’s species-abundance curve (i.e., sharp decrease in abundance and breadth coverage). In contrast, abundant MAGs were defined as those showing >0.1% relative abundance and coverage breadth more than 90%. Note that robust detection of a MAG and estimation of its relative abundance is achieved with breadth 10% or higher [67]. Hence, MAGs are reliably detectable at 0.1% relative abundance or somewhat lower, and the 0.1% abundance threshold is not a mere artifact of sequencing effort applied (e.g., inability to detect a MAG). Our definition seems to largely agree with the literature on defining rare biosphere as well [68].

**Figure 2.**
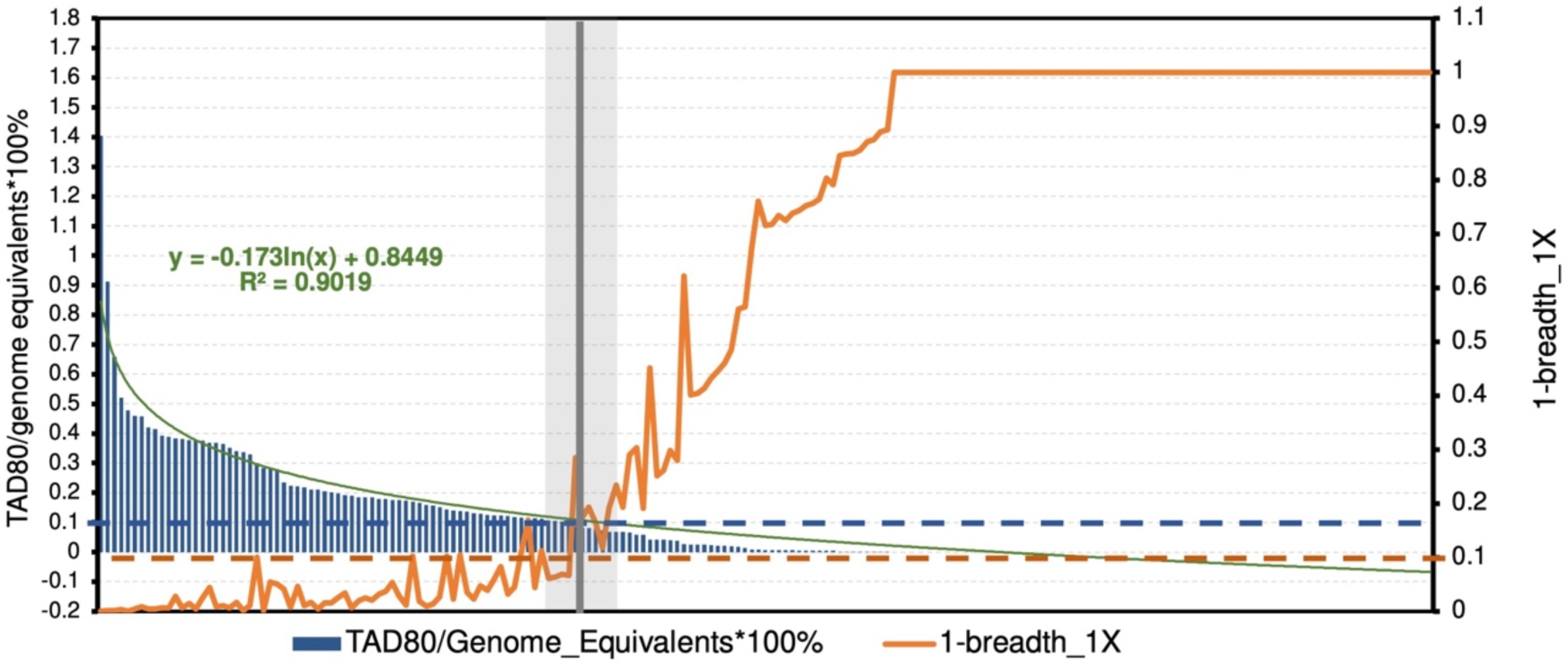
Our approach to define abundance vs. rare MAGs. MAG coverage depth (left y axis, blue bar) and coverage breadth (right y axis, orange line, shown as 1-coverage breadth) distribution for one metagenomic sample (pico127). Thus, X axis is MAG rank by abundance, estimated as coverage depth (i.e., TAD80 values normalized by genome equivalents). Dashed blue and orange line represent normalized coverage depth 0.1% and coverage breadth 0.1, respectively. Grey area and vertical line (center of area in terms of x axis) indicate regions where both coverage depth and breadth drop sharply as abundance rank increases. The vertical line was therefore used to define abundant (MAGs to the left of the line) vs. rare (MAGs to the right of the line) MAGs. Green line is a log fitting of coverage depth vs rank with fitting function shown above the line. For detailed model fitting of coverage depth distribution, see Supplementary figure S7.

### Testing the insurance hypothesis: rare MAGs often contribute to community response

We first tested for the predicted signature of the insurance hypothesis in each of the six major disturbance events sampled by our time series metagenomic datasets. Since our metagenomic sampling is part of a much longer (3-year) 16S amplicon-based time series dataset, we selected samples for metagenome sequencing to represent the pre-disturbance according to the 16S diversity and composition relative to additional samples available in the time series for the same season that the disturbance event took place.

That is, these samples did not have a significant difference compared to the previous week, and thus served as useful controls for assessing the effects of the disturbance [69]. We first checked the overall MAG relative abundance changes across all disturbance events and found that for each event, many MAGs showed clear changes when comparing disturbed samples with pre-disturbed samples (Figure S9). We found that for all 6 disturbance events, there were between 2 to 22 microbial populations represented by MAGs (Figure 3) that switch from being rare to being abundant during the event based on the criteria mentioned above to define rare MAGs. Further, these MAGs represented between 0.5% to 6.9% of the total community based on the number of reads mapping on the MAGs (relative to the total reads of the sample), depending on the event considered (Figure S10). There were also 8 to 29 MAGs that switched from being abundant to rare, representing 0.4% to 1.8% of the total in the corresponding (disturbed) sample. For each event examined, there were always more MAGs assigned to the abundant-to-rare category than to the rare-to-abundant category, except for event 8 (spr13_warmer_8) (Figure 3). Further, the fraction of reads mapping to rare-to-abundant MAGs was higher than that of abundant-to-rare MAGs in the disturbed samples in three events out of total six (Figure S10). Note that about 5.1% to 18.2% of the total reads in a sample were mapped to all available MAGs (not only MAGs in the 3 categories above but also MAGs assigned to rare-to-rare category), which represents a substantial fraction of the microbial communities sampled; much higher fraction than previous, isolate- or lab-based approaches to study similar questions were able to assess.

**Figure 3.**
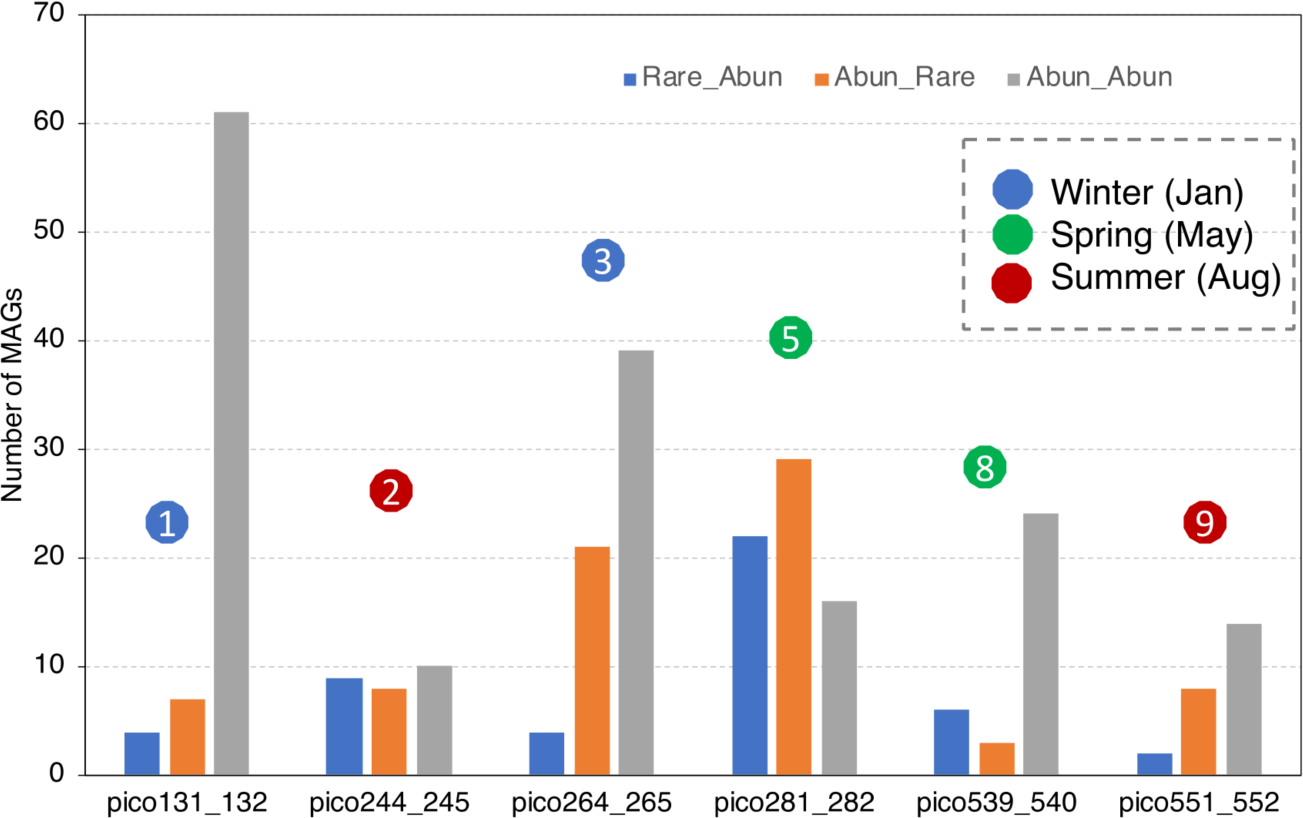
Responses of abundant vs. rare MAGs to the disturbance events. The figure shows the number of MAGs for each of the three categories assessed: abundant MAGs that remained abundant after the event, MAGs becoming abundant from rare, and MAGs becoming rare from abundant for each of the disturbance event. For one given event, if MAG’s relative abundance and coverage breath fall below the threshold of being abundant in the pre-disturbance sample and fall above the threshold of abundant in the disturbed sample, this MAG will be in the category of Rare_Abun. Similar rules applied for other two categories. Disturbance events are labelled with a number as in Figure 1 (**see Figure S1 for details**). See Figure S10 for total relative abundance of MAGs for each category.

For event #5 representing an extreme warm week during the spring time (spr12_warming_5), there were 21 rare MAGs that became abundant, while 29 abundant MAGs switched to rare. Sixteen abundant MAGs remained abundant during this event, representing 4.5% of the total community. The total relative abundance of MAGs that switched from rare to abundant was around 7% while the abundant MAGs that switched to rare made up about 1% of the community in the disturbed sample (but was 5.3% before the disturbance; Figure S10). Further analysis showed that the rare MAGs that became abundant during this warming event (spr12_warming_5) encode more metabolic pathways related to carbohydrates degradation and fewer pathways related photosynthesis compared to abundant-to-rare MAGs (Figure S16). These results suggested that there were more carbon compounds in the surface seawater during this warming event compared to pre-disturbance sample, and microbial populations that were more efficient in utilizing those carbon gained a competitive advantage over those that lacked these pathways (e.g., abundant to rare MAGs). Amplicon based analysis also showed consistent results: a “spring bloom” indicating overturn in the phytoplankton was obvious based on the amplicon datasets [22]. However, two weeks after the event, 14 of the MAGs that had become abundant from rare during the event became rare again (77.8% of the category) while 11 MAGs that became rare from abundant became abundant again (66.7% of the category), revealing that the sampled microbial communities were resilient. These results reveal that the rare populations provided insurance for ecosystem functioning when undergoing a short-term, strong disturbance event. That said, the importance of abundant populations that remained abundant cannot be underemphasized because the latter MAGs represented a higher fraction of the total community than rare-to-abundant MAGs in all events, including the disturbance event 5 (spr12_warming_5) mentioned above (Figure S10).

There are only three disturbance events with available post-disturbance samples: the other two being win11_storm_2 and win12_warming_3. For win12_warming_3, we observed a relatively small number of MAGs changing category, i.e., becoming rare-to-abundant or abundant-to-rare (25% and 33.3% of the total MAGs in each category, respectively) compared to the warming event mentioned above. For win11_storm_2, 44.4% of MAGs became rare-to-abundant and 25% of MAGs became abundant-to-rare, and all these MAGs recovered to their pre-disturbance level (category) after the event. For the remaining disturbance events, we observed 50% (out of 4 MAGs in total) and 42.8% (out of 7 MAGs in total) of the rare-to-abundant and abundant-to-rare categories to recover to pre-disturbance levels for the win11_warming_1 event, respectively, while 50% (out of 2 MAGs) and 25% (out of 8 MAGs in total) of the same categories recovered post-disturbance for the sum13_storm_9. Overall, except for event win12_warming_3, about half or more of the taxa that represented rare-to-abundant populations recovered after the event (that is, returned to rarity), further indicating that while rare populations play a role in community resilience by responding to niches created by the disturbance, most of these taxa return to being rare post-disturbance. However, except for spr12_warming_5, less than a half of the abundant-to-rare populations recovered spr12_warming_5 to become abundant again, indicating that rare and abundant populations may respond differently to disturbances.

### Testing the disturbance-specialization hypothesis: generalists are favored by disturbance

We additionally examined the types of organisms that responded to disturbance in term of generalist vs. specialist lifestyle. For the latter, we used the proportion similarity index that measures population environmental preference by examining abundance changes across categories of available samples (PS index) (Table S3). We categorized the upper and bottom third of MAGs based on ranked PS index (see Materials and Methods for details) as generalists and specialists, respectively. Notably, our ranked based definition of generalists is generally consistent with the Levin’s Breadth Index and PS index implementation in MicroNiche (Figure S19), but it is more appropriate for the data available here as explained in the Methods section. For rare populations that became abundant after the disturbance, a larger fraction of them were generalists (60% to 78%) than specialists (16% to 24%) except for the first winter disturbance, which was in fact a resilience event (community recovered to pre-disturbance) (Figure 4). On the other hand, for abundant populations that became rare, most of them were specialists, except for sum11_cyclone_2 and sum13_cyclone_9, which were summer cyclone disturbance events (Figure S11). Functional gene content analysis revealed that generalists had smaller genome sizes and more compacted genomes than specialists (Figure S12), and a slightly higher fraction of extracellular function proteins encoded in their genomes (Figure S13). These results support the disturbance-specialization hypothesis to explain, at least partially, responses to nearly all disturbance events.

**Figure 4.**
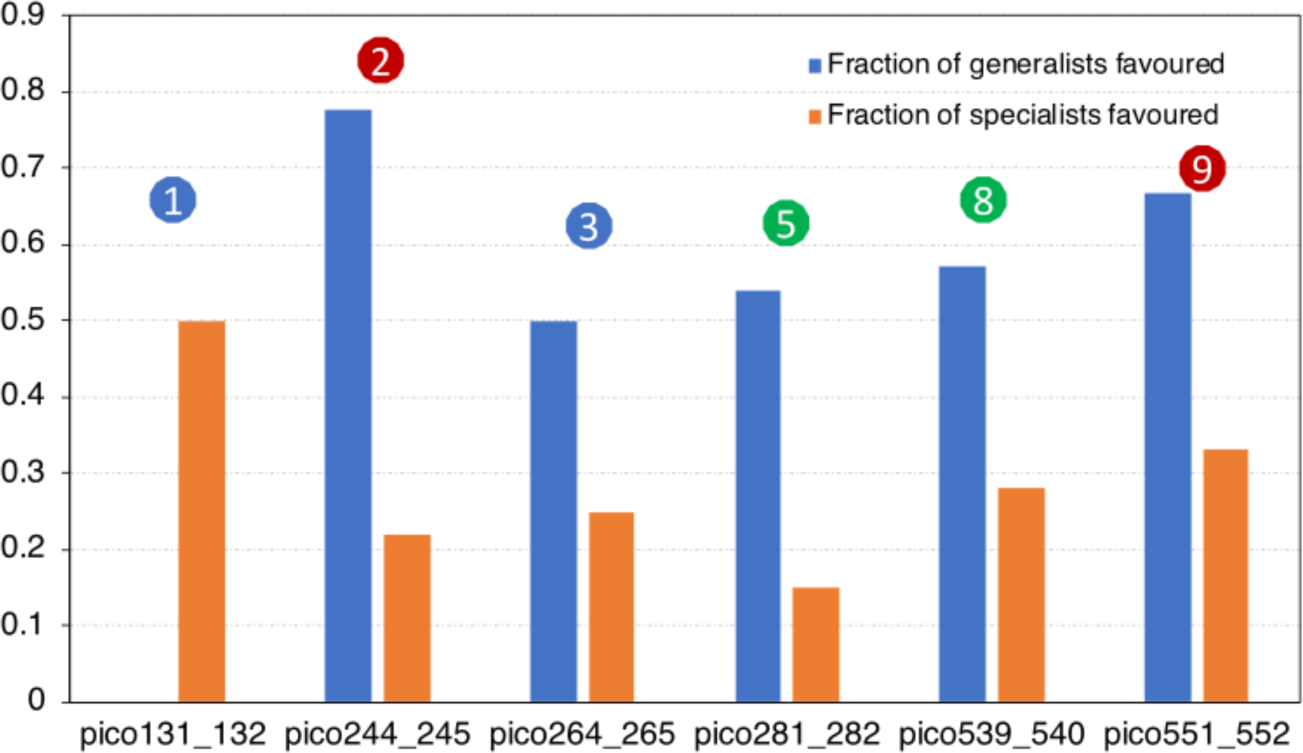
Fraction of specialists vs. generalists selected by each disturbance event in terms of number of MAGs. Selected MAGs are those that became abundant from rare in each disturbance event shown in Figure 3. Disturbance events are labelled with a number (see Figure S1 for details).

For winter (warming) disturbances (win11_warming_1 and win12_warming_3), the populations that remain abundant were identified mostly as specialists and only a small fraction of them were generalists. In contrast, for summer disturbances (sum11_storm_2 and sum13_storm_9), most MAGs that remained abundant were generalists (6 and 9 respectively), suggesting that these generalists could survive strong water body mixing disturbances caused by cyclones. For spr12_warming_5, the minority of abundant MAGs that remained abundant before and during the event were identified as generalists (4 out of 11), contrasting with spr13_warming_8, in which the majority of abundant MAGs that remained abundant before and during the event were generalists (17 out of 21). These patterns suggested that abundant populations of the community, which could be either generalists or specialists, are generally resistant to the effects of the disturbance events studied, consistent with the results reported above at the whole-community and MAG levels.

## Discussion

Quantifying the importance of rare populations for microbial community response to disturbances represents a cornerstone question in microbial ecology and is key to testing ecological theory as well as modeling the effects of global change on ecosystem function and diversity. We observed that a large fraction of the detected MAGs remained abundant before, during and after disturbance (resistant) for all the disturbance events studied (Figure 3, Figure S10), revealing that these genomospecies are insensitive to short term disturbances and provide stability for the ecosystem. We also found that rare prokaryotic populations became abundant after the disturbance event and made up between 0.3% and 7% of the total community, depending on the event considered. However, more than half of these populations returned to rarity one or two weeks after each disturbance, consistent with the concept of conditionally rare taxa (CRT) defined by Shade and colleagues [7] and also the results of amplicon based analysis of same samples [22]. Such conditionally rare taxa may therefore provide resilience to the system undergoing disturbances and our approach here quantified this contribution in terms of relative abundance of the total community. As community assembly theory also predicts, CRT that are deterministically assembled (that is, selected by disturbance conditions) may contribute to the ecosystem processes/functions as much as the abundant taxa do [3]. In agreement with these interpretations, we also found that rare-to-abundant population genomes harbored more metabolic pathways related to carbohydrate degradation compared to abundant-to-rare populations, the latter were clearly disfavored by the spr12_warming_5 (spring turnover blooms) disturbance events. It should be mentioned, however, that this pattern was clear only in one of the two warming events (spr12_warming_5; the other being spr13_warming_8) and despite the shared characteristics between the two events such as the increase of chlorophyll A concentration, an indicator of primary production.

The lack of universal patterns among the disturbance events is likely due, at least in part, to the fact that the disturbance events were all largely distinct from one another, even for similar winter “warming” disturbances, and each event favored different MAGs and functional traits (so high inter-event diversity). For example, for the two summer hurricane disturbance events, we did not observe consistent rare-to-abundant population dynamic patterns (e.g., the same genomospecies did not show similar changes), partially because these cyclone events caused the strong mixing of both costal sediment and seawater. We also saw different patterns in the warming winter (win11_warm_1) and rainy winter (win12_rainy_3) disturbances, which, nonetheless, was not surprising because these two disturbance events were characterized by different nutrient changes such as ammonium concentrations (Table S1, pico131 and pico256 for 2 events respectively). Those inconsistent responses, in general, are somewhat expected because capturing natural disturbance events and associated metadata is challenging from the perspective of sampling. Although replicating disturbances is not possible (e.g., all storms are not identical), (biological) replicate samples are possible, and we will try to obtain such replicate filters in future studies. Also, a lower number and quality of MAGs were recovered for the datasets representing the second hurricane event due to relatively lower sequencing coverage (Figure S2, sample pico551 and pico552 were 1/3 of other samples in terms of size), which somewhat limited our resolution into the population dynamic patterns. Deeper sequencing efforts could further facilitate the study of patterns of rare population dynamics.

How to define rare populations or species has been a challenging task, and arbitrary thresholds based on relative abundance of 16S rRNA gene sequences (e.g., 0.1%) have been commonly used [15]. Here we defined rare populations or species based on the species-abundance curve that was derived from the sequencing depth and breadth coverage of recovered MAGs (normalized by total genome equivalents), while considering the limit of detection of our metagenomic sequencing effort. Specifically, the TAD80 metric used for calculating abundance was larger than the genome breadth coverage that corresponded to the limit of detection, which was defined as - ln(0.9)/genome equivalents [70]. Therefore, the commonly derived 0.1% threshold for rare taxa by our analysis, and our definition of rare MAGs, were robust. Although our derived threshold often matched the previous (arbitrary) thresholds, we anticipate that the threshold may differ for other samples and/or environments. Hence, the approach outlined here based on species abundance curve should be useful for future studies. Further, the TAD80 (coverage depth), a metric used to define population abundance, can remove highly conserved regions and regions recently subjected to horizontal gene transfer. If those regions are not removed, the sequencing depth, and thus abundance can be overestimated. TAD80 can also remove regions with high gene-content micro-diversity, which tend to underestimate depth. Thus, TAD80 provides reliable estimations of relative abundance and thus, species abundance curve [57,71]. It should be mentioned, however, that for the species abundance curves we did not observe a clear inflection point that could be used to define rare species in a more natural and objective manner compared to the use of a predetermined threshold (e.g., 0.1% abundance). Instead, the curve often appeared to be a monotonic, log-normal decrease with no obvious inflection points but with a sharp change in slope. Hence, the threshold to define rare taxa may appear somewhat arbitrary even with a species abundance curve available, although we do recommend the TAD-80-based methodology outlined above as a more robust and well-defined approach that takes into account (normalizes for) different sequencing efforts between samples (see also below).

Rare taxa have been defined based on several different approaches. For instance, Debroas et al. defined rare biosphere as unassembled reads with low sequence depth coverage, without obtaining population genomes, and bioinformatically annotated those unassembled reads by mapping them to pre-annotated databases in order to infer functions carried by the rare taxa [72]. However, this approach cannot link these functions with specific taxa and derive species abundance dynamics since these unassembled short reads are typically not linked to each other and/or a phylogenetic marker. Our approach to define rare population based on MAG abundances can link taxa with their functions, providing an important advantage over previous literature, albeit the number of taxa (or MAGs) that we were able to study was relatively small (e.g., the MAG has to be abundant enough in at least one sample of the series to be recoverable by sequencing). Further, binning rare population MAGs is subjected to artifacts (more so than for abundant MAGs) because such populations show low sequencing coverage, and thus are typically represented by short contigs that are not easy to bin (Figure S14, white and orange datapoints) [73,74]. Long read metagenome sequencing could improve assembly and binning even of low coverage MAGs, especially in combination with new binning algorithms, such as those employing deep variational auto-encoder [75] and graph-based embedding method [76] or both [77].

To further understand the key functional and/or ecological differences between taxa that responded to disturbances by changes in their abundance relative to those that did not, we tested the disturbance-specialization hypothesis that generalists are favored while specialists are disfavored by disturbance. Our results provided some support for this hypothesis based on the PS index, which is a direct way to measure taxon relative abundance changes across the entire dataset to define generalists vs. specialists. Our conclusion that more generalists taxa than specialists are favored by most disturbance events studied here is consistence with those of recent studies based on laboratory mesocosms [78]. Specifically, Chen and colleagues concluded that generalist taxa are more metabolically flexible. Consistently, we found that generalist have -on average- a smaller and more compacted genomes, which probably provides an (more) efficient metabolic strategy and a selective advantage during changing conditions. The smaller genome size is also consistent with the Black Queen Hypothesis that has been used to explain the ecological success of small genomes [79]. Further, we found that generalists have a slightly higher fraction of their genome devoted to extracellular protein functions compared to specialists, which could be another reason why generalist were favored by the disturbance events. That is, extracellular proteins represent mostly enzymes that degrade or transport public goods that presumably remain abundant throughout the disturbance.

It should be noted, however, that there is no standard or universally accepted definition for generalist/specialist except probably for Levin’s niche breadth index [62,80] and Proportional Similarity Index [63]. We preferably employed the PS index because it allows abundant and widespread species to be also classified as habitat specialists when they show relatively high abundance in a single category of samples (so prevalence in more samples matters in addition to their abundance or absolute abundance) [57,64]. We obtained largely consistent results to those reported above with the PS index when we used the PS implementation in MicroNiche package, as well as with Levin’s niche breadth definition (Figure S19)[57]. Further, the PS index, based on absolute abundance data, is commonly used in macroecology compared to Levin’s definition [64]. We used relative abundance of microbial taxa as compared to the absolute abundance typically used in macroecology since absolute abundances were not available for our datasets. Performing this type of analysis with absolute abundances and a more detailed measurement of nutrient availability in each sample in future studies, could further corroborate the conclusions presented here.

It should also be noted, however, that the definition of rare populations used could affect our findings and conclusions. For example, the sequencing effort applied presumably affects the limit of detection established, and thus our definition of rare taxa. Under-sampling (in term of sequencing coverage) of microbial communities, especially in highly complex environmental settings, leads to poor assembly and MAG recovery. Further, this limitation could also affect the species abundance curve and thus, the threshold for rare taxa derived from the curve. For example, for non-subsampled reads, we obtained a total of 412 dereplicated MAGs compared to 198 when we subsampled the datasets to provide similar sequencing effort across datasets, further indicating that more MAGs can be obtained with a higher sequencing effort. More available MAGs could somewhat affect the species abundance distribution curve, and thus our empirical definition of rare population changes (e.g., threshold to be mostly around 0.05% instead of the 0.1% used) (Figure S17). Sequencing coverage could also affect the PS index because with more sequencing effort some undetected genomes in under-sampled samples could become detectable. Nonetheless, we expect that this sequence coverage limitation applies evenly to all samples and rare taxa, and thus should not significantly affect our PS index values for most taxa and our derived conclusions. It would be valuable to see if our conclusions would change with higher sequencing coverage and absolute abundance measurements of each species in future studies of our sampling and other sites.

Our work also showed that natural microbial community are remarkably resilient at the whole community level. Both diversity (Figure 1c and Figure S4 a and b, MAG recovery results section) and community composition (Figure S5) recovered after disturbance, consistent with previous findings in a lake ecosystem and elsewhere [81]. Metagenomic-based functional diversity analysis showed that functional redundancy could play a key role in this resilience because we observed a decoupling between taxonomic diversity changes and functional diversity changes (Figure S6), which presumably reflects functional redundancy among the taxa that changed in abundance [82]. The recovery of the rare-to-abundant populations also supported the strong resilience of the sampled communities (that is, going back to being rare after disturbance) even though these populations typically accounted for a smaller fraction of the community typically, i.e., 1-7% of total. The fact that we did not observe high similarity in the microbial community responses to similar disturbance events in the same season underscores the great diversity of microbial communities as well as our inability to measure the key parameters changes by each event e.g., diversity of organic compounds released or exact physicochemical properties that may slightly differ among similar disturbance events. Year to year variation, e.g., stochastic birth/death of microbial populations, could also be another reason we did not see consistent responses among similar events [83]. Multiple samples that represent the prevailing, not-disturbance-associated, microbial communities in each season would be needed to quantify the effect of stochastic processes on the results reported here. For instance, it is possible that some of the MAGs that are reported above to change from the abundant to the rare categories during the event (or vice versa) might represent such stochastic processes-as opposed to deterministic processes caused by the event. We believe that the effect of such stochastic processes on our results is limited because in a couple samples that the post-disturbance sample resembled closely the pre-disturbance sample in terms of 16S-based microbial community composition, and thus allow us to assess stochastic processes, such as for win12_rainy_3 (see Fig. S4), the MAG abundances were much more similar between the pre- and post-disturbance samples relative to the disturbance sample (Fig. S18). Therefore, most of the MAGs reported to change between the abundant and rare categories are presumably due to the effects of the events rather than stochastic processes.

In conclusion, we found microbial communities in the coastal ocean of Southeast USA to be both resistant and resilience (depending on whether the corresponding populations were abundant and stayed abundant or became rare during the event and bounced back to be abundant after the event, respectively) against natural disturbances, and provided evidence in support of the disturbance-specialization and the insurance hypotheses. Further, we provided a new approach based on the species abundance curve to define rare vs abundant populations as well as generalists versus specialists based on a time-series metagenomic sampling. These definitions and approaches could be helpful for future ecological studies aiming at answering questions related to the role and importance of the rare biosphere. Collectively, the findings presented here advance our understanding of natural microbial community response to disturbance, and thus, could be useful for ecosystem management, e.g., microbiome rescue [8] in a changing world.

## Data Availability

Raw metagenomic reads, amplicon reads can be found at NCBI, under accession number PRJNA803723.

## Author Contribution and Funding

J.Z, L.M.R and K.T.K designed the work. Z.W and D.H did the sampling and environmental analysis. J.Z and G.B did the bioinformatic analysis. J.Z and K.T.K wrote the manuscript. This work was supported, in part, by the US National Science Foundation to D.E.H. (OCE 1416665) and to K.T.K. (OCE 1416673 and DEB 1831582).

## Acknowledgment

We want to thank PACE at Georgia Tech for providing computational resources.

## Supplementary Figures

**Figure S1.**
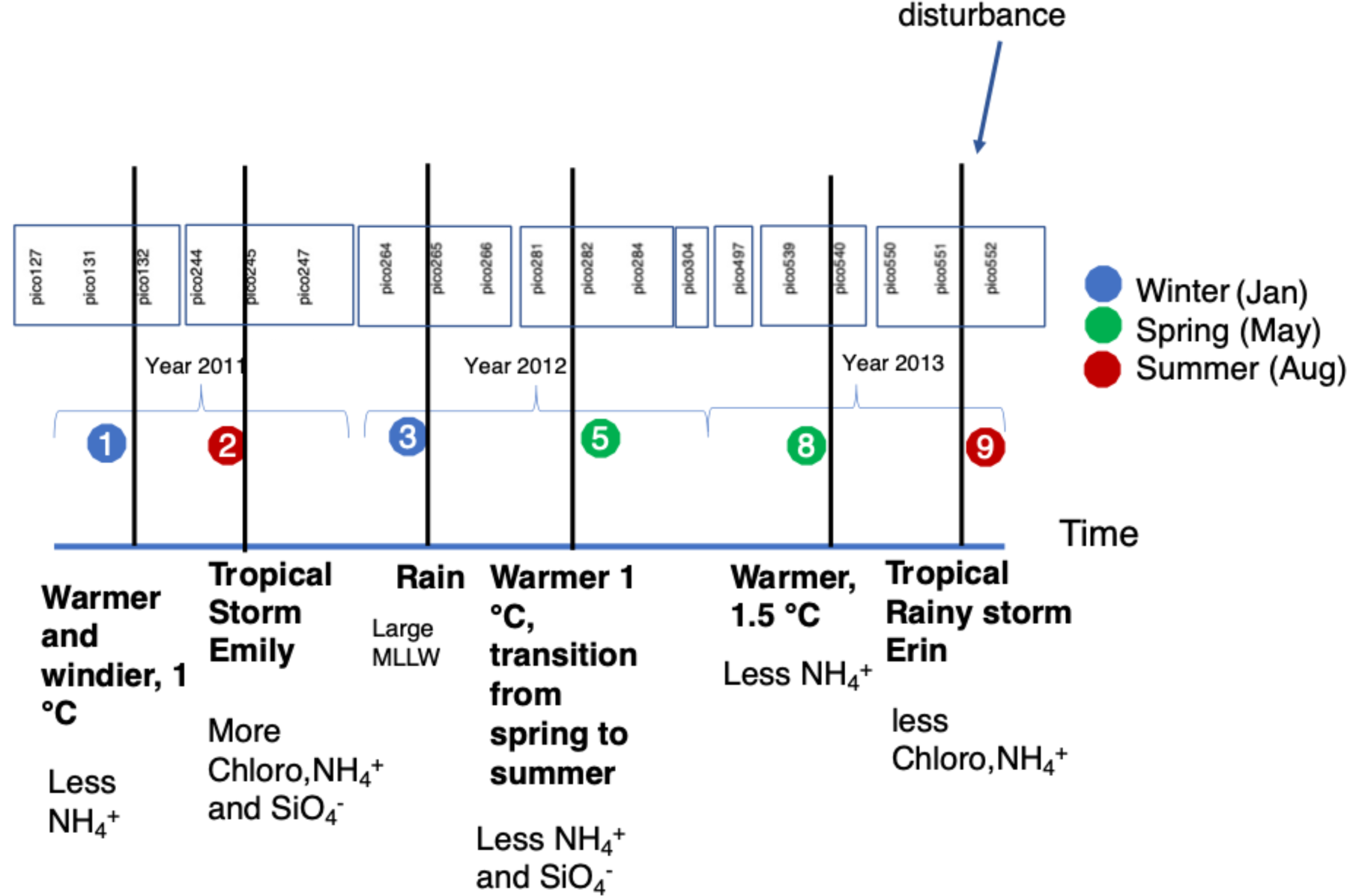
Details of the Pivers Island Coastal Observatory (PICO) time series samples. Sample names are labeled chronologically within boxes, the color of which corresponds to seasonality along the three-year sampling period. Continuous numbers in the name of samples indicate samples taken one week apart (e.g., pico281 and pioc282 are sampled from 2 adjacent weeks, while pico284 is sampled 2 weeks after pico282). Each vertical line indicates a disturbance event and is labelled with a number for convenience. Description below each vertical line shows the details of each disturbance and the most pronounced differences in environmental parameters measured. MLLW stands for the average lowest of the two low tides of a week.

**Figure S2.**
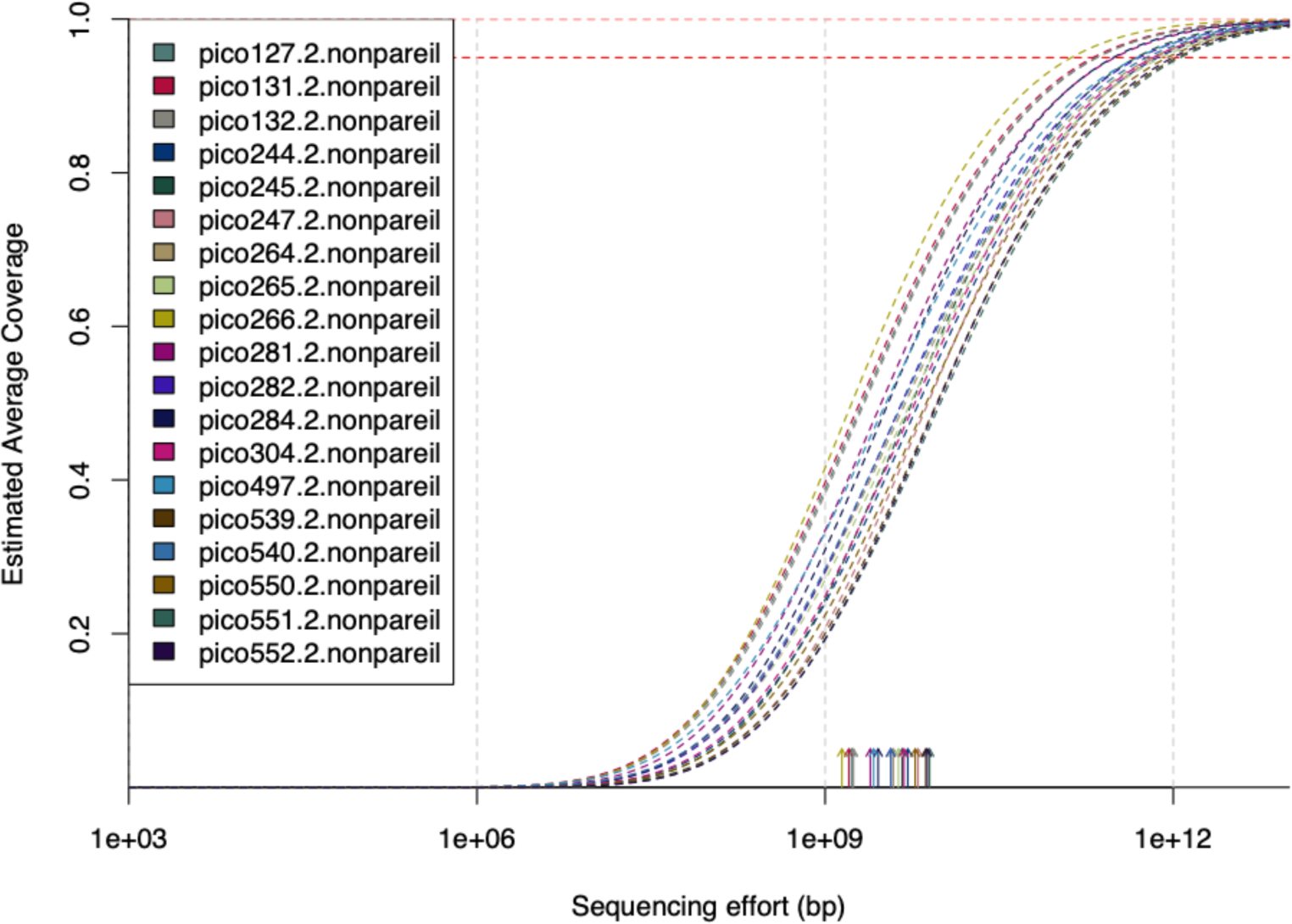
Nonpareil curves showing the coverage of subsampled metagenomes. Solid lines show the estimated average coverage (y-axis) as sequencing efforts increases (x-axis); dashed lines represent the projections for 95% and 99% coverage (horizontal dashed lines on the top). Only reverse reads were used for coverage estimation according to Nonpareil author recommendation for sequences used to not be linked/associated to each other (independent observations); forward reads showed similar curves for each sample (not shown).

**Figure S3.**
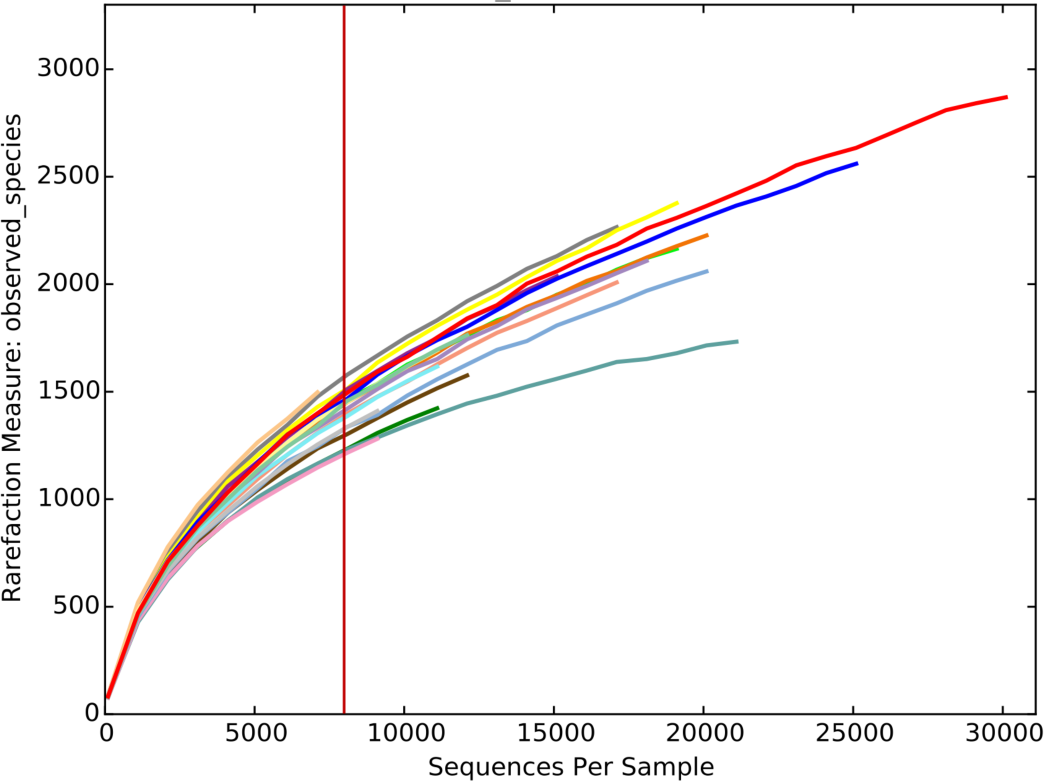
Rarefaction curve for extracted 16S-carrying reads from metagenomes. Each curve represents a sample. The vertical red line shows the number of reads subsampled (8000) for downstream analysis.

**Figure S4.**
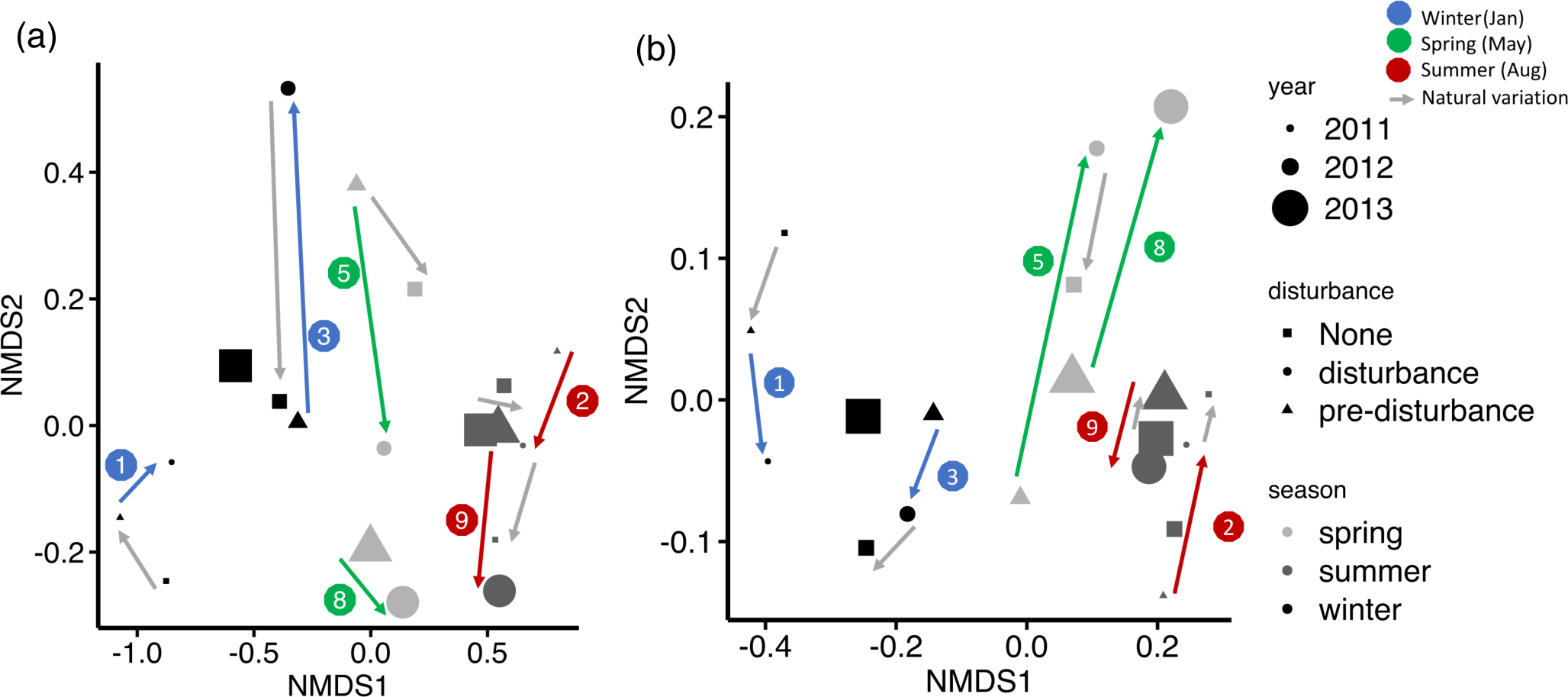
NMDS plot of amplicon 16S rRNA gene sequences (a) and extracted 16S rRNA gene-carrying reads from metagenomes (b). Arrows show the direction that the microbial community composition changed by each disturbance event. Grey arrows show the natural variation of metagenomic composition.

**Figure S5.**
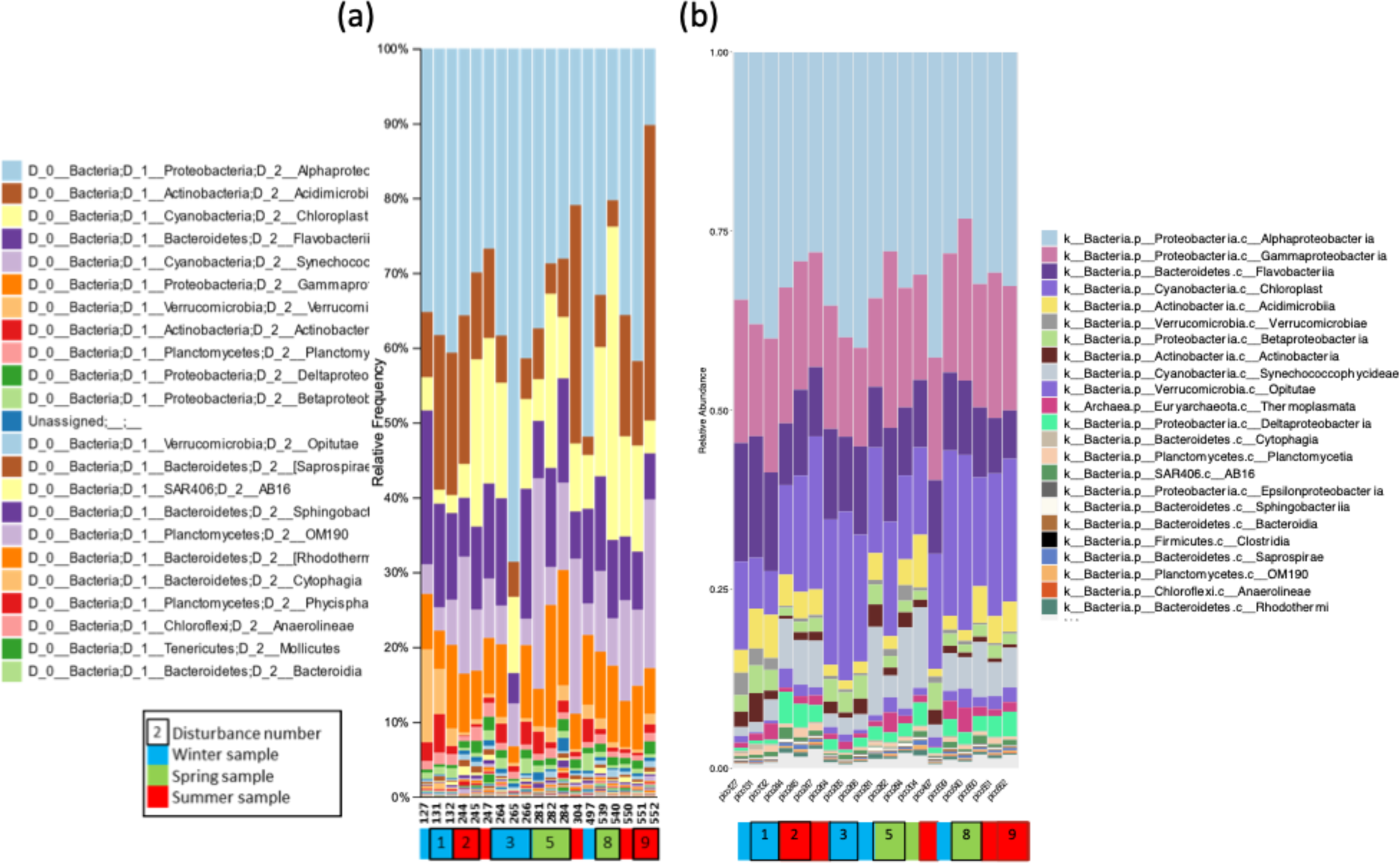
Class level microbial community composition (relative abundance) for amplicon 16S (a) and 16S-carrying reads extracted from metagenome (b). Each column represents a sample, with sample details provided by the color box. Disturbance events are labelled by the same number as in Figure S11. See Figure S1 for detailed explanation for each disturbance event.

**Figure S6.**
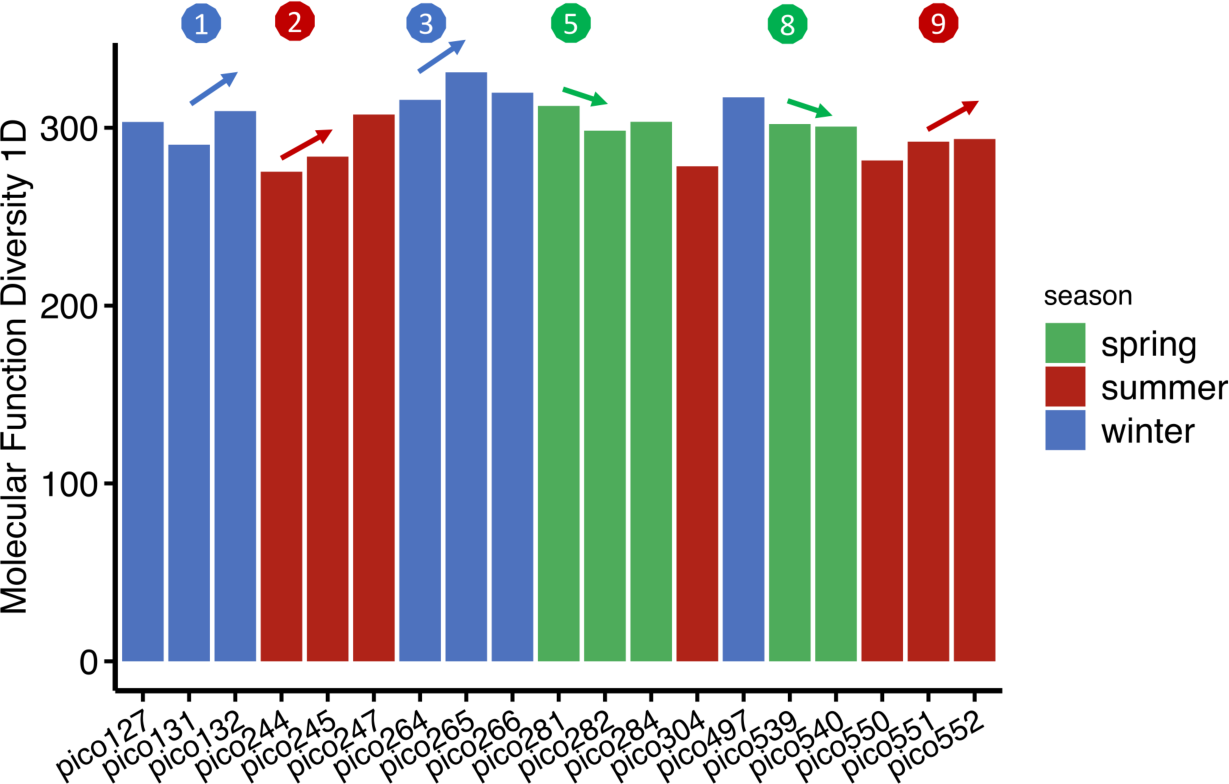
Changes in molecular functional diversity of metagenomes by each disturbance event as revealed by mapping reads to annotated functional pathways (See Methods & Materials).

**Figure S7.**
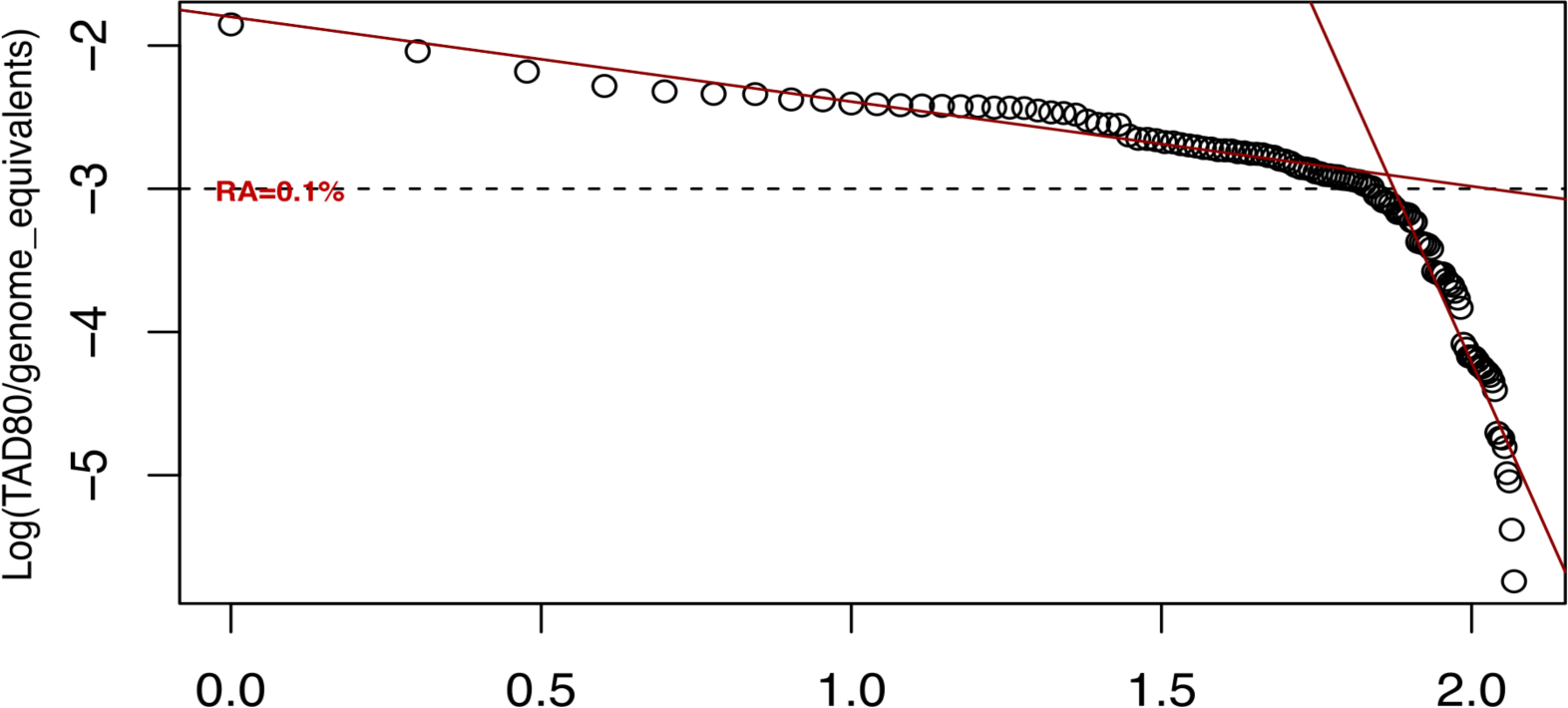
Log-log fitting of normalized sequence coverage depth vs. abundance rank for all MAGs used in the study. Two linear fittings were performed using MAGs with normalized coverage depth larger than 0.1% and smaller than 0.1% respectively (R^2^ > 0.7). A clear difference in slop of two fit lines indicate a sharp decrease for normalized coverage depth around 0.1%.

**Figure S8.**
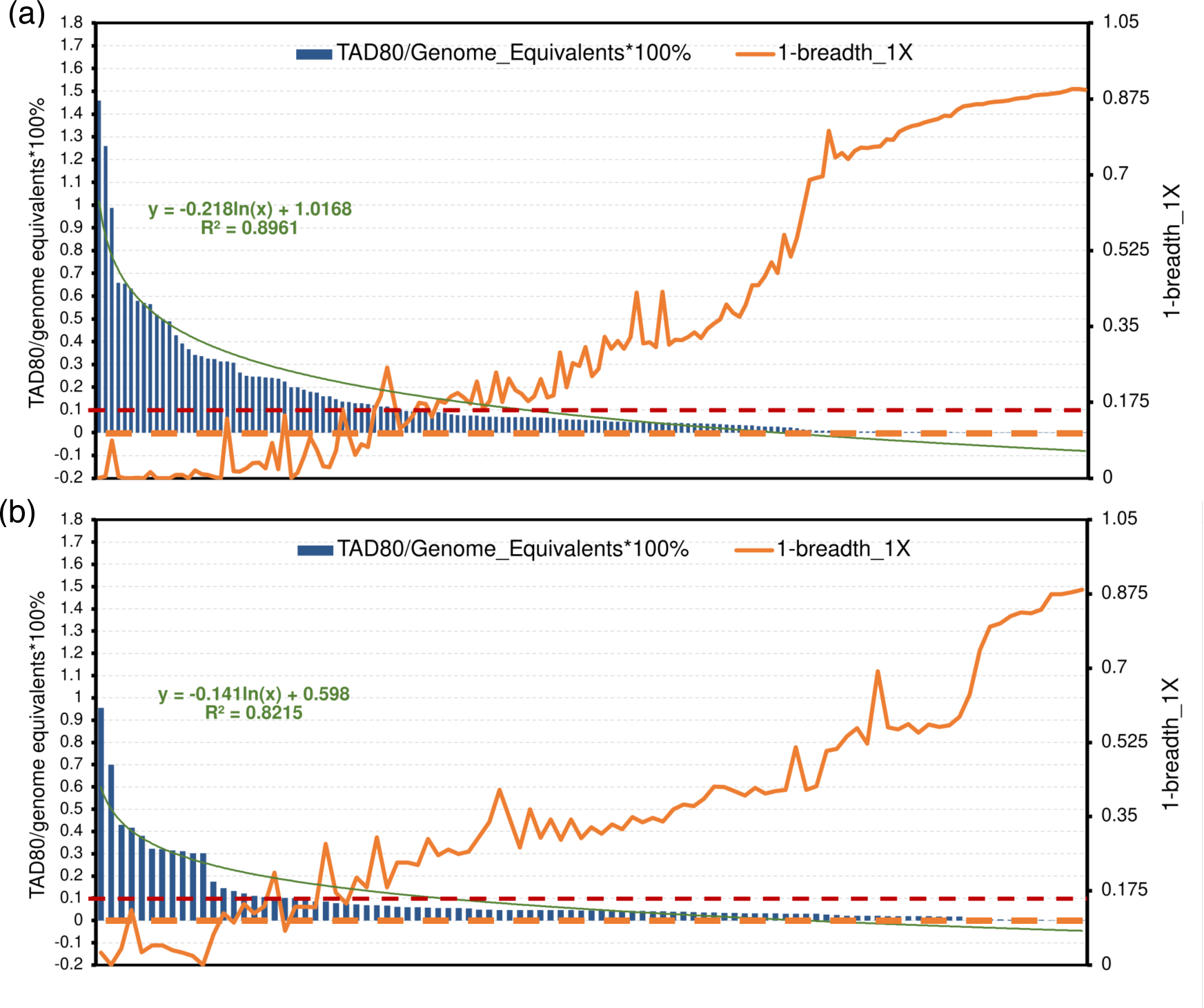
MAG coverage depth (left y axis, blue bar) and coverage breadth (right y axis, orange line, shown as 1-coverage breadth) distribution for two metagenomic samples, pico284 (a) and pico247 (b). This figure shows two additional examples and consistent patterns to those observed in Figure 3. X axis is MAG abundance rank based on coverage depth (TAD80 normalized by genome equivalents). Dashed red and orange lines represent normalized coverage depth 0.1% and coverage breadth 0.1, respectively. Green line is a log fitting of coverage depth vs rank with the corresponding function shown above it. Before subsampling, pico284 has similar sequencing depth with pico127 while pico247 is the shallowest sequenced sample of the three, thus only a few MAGs show high coverage depth (i.e., less reads could be mapped to the dereplicated 198 MAGs).

**Figure S9.**
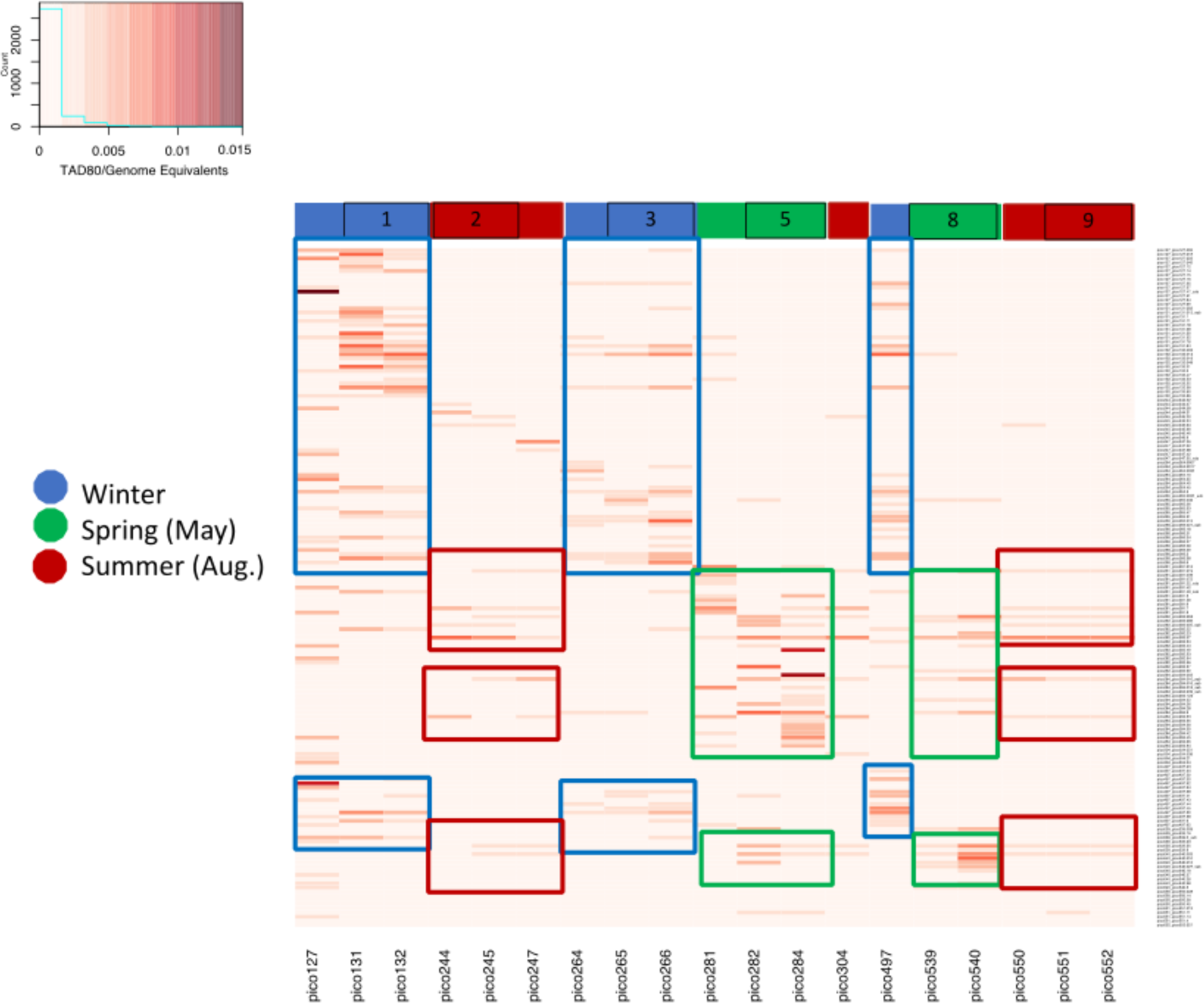
Heatmap of MAG relative abundance. Each row represents a MAG while each column represents a sample (see key for sample designation by color). Disturbance events are labelled by a number as in Figure 1. See Figure S1 for detailed explanation for each disturbance event.

**Figure S10.**
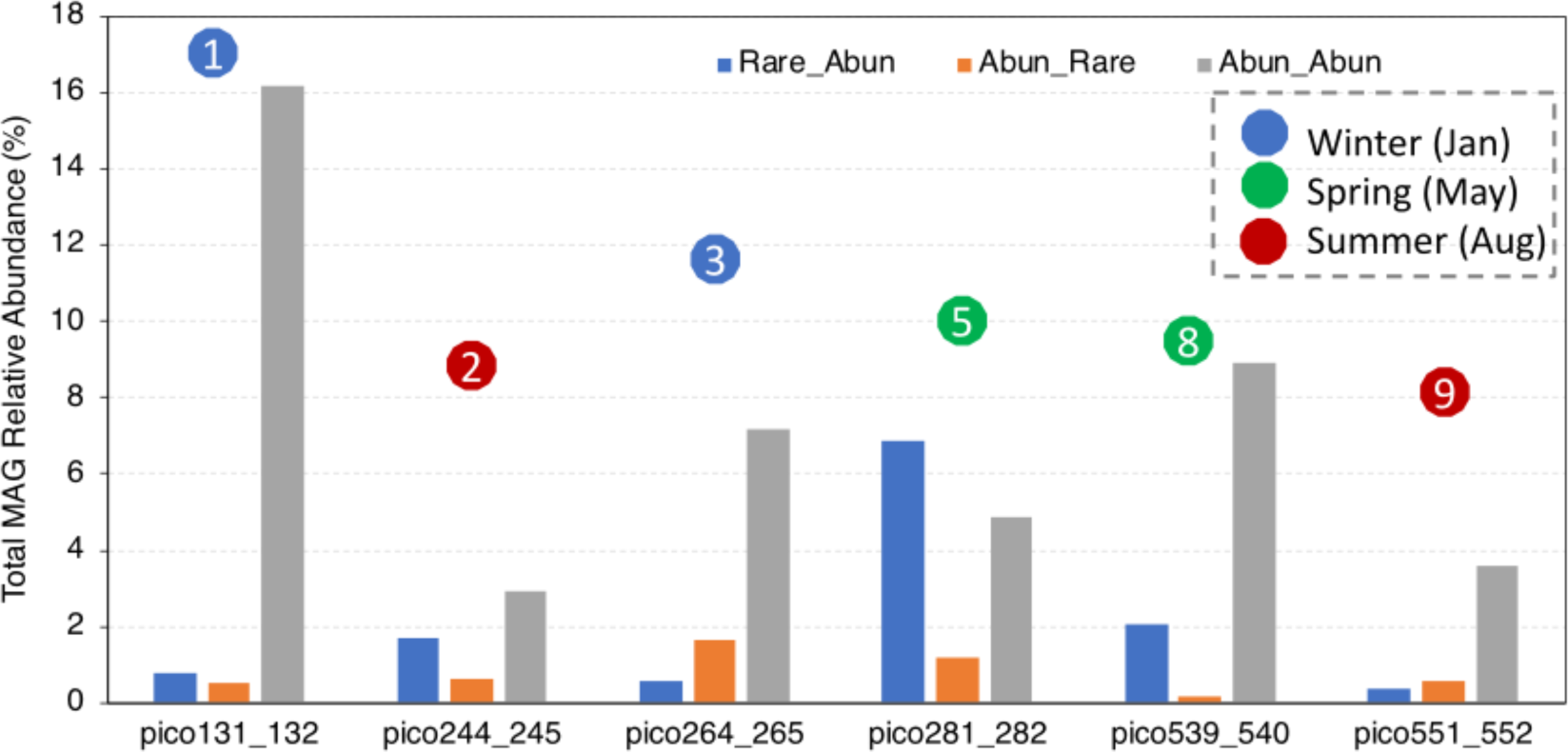
Total relative abundance of MAGs assigned to each category of abundance change. Categories were: remain abundant, become abundant from rare and become rare from abundant for each of the disturbance event (comparing samples before and after each disturbance event). Disturbance events are labelled with number (see Figure S1 for details).

**Figure S11.**
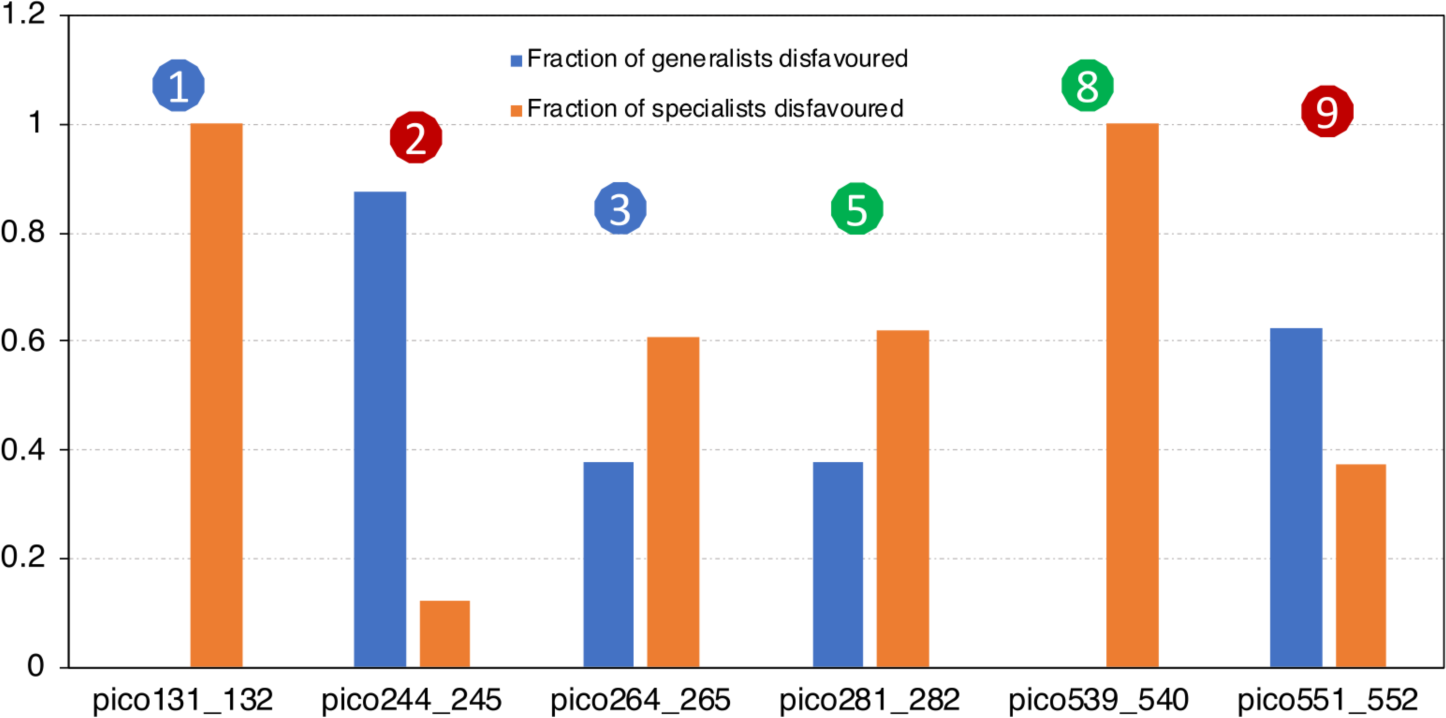
Fraction of specialists and generalists disfavored by each disturbance event. Disfavored MAGs are those that become rare from abundant by each disturbance event (similar to Figure 4 but the opposite pattern). Disturbance events are labelled with number (see Figure S1 for details).

**Figure S12.**
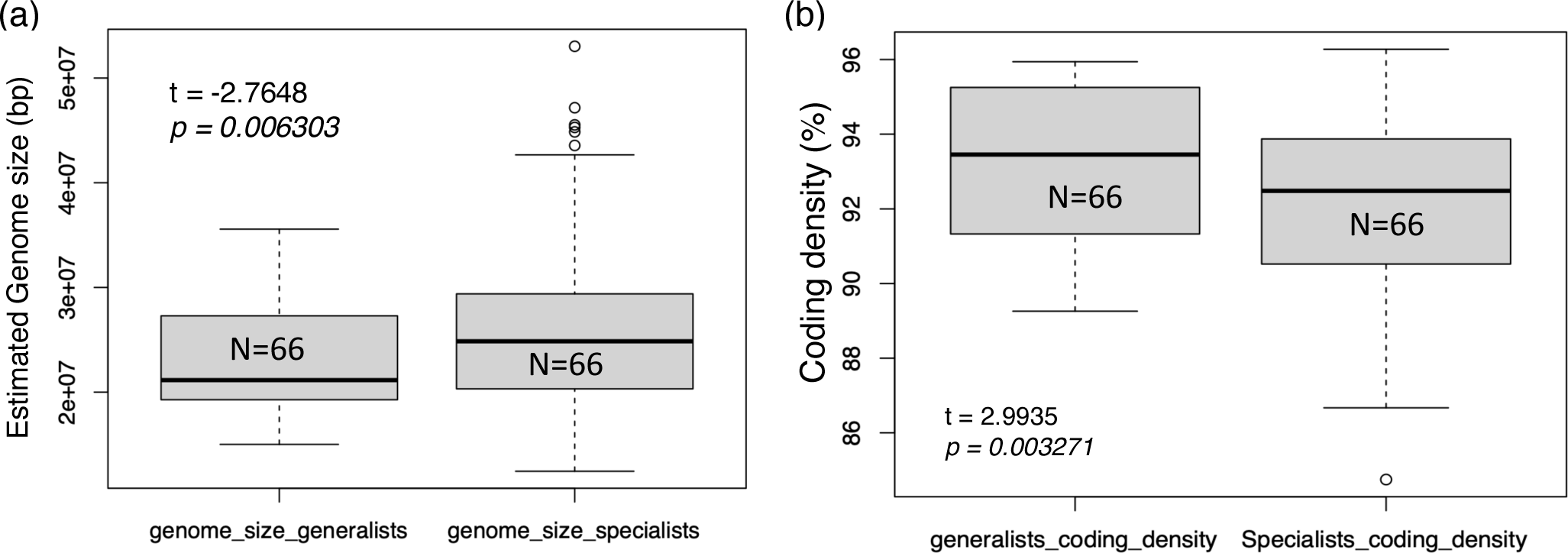
Estimated genome size (a) and coding density (b) differences between generalists and specialists. Both T-test and Mann–Whitney test were significant (note the p-values shown on the graphs).

**Figure S13.**
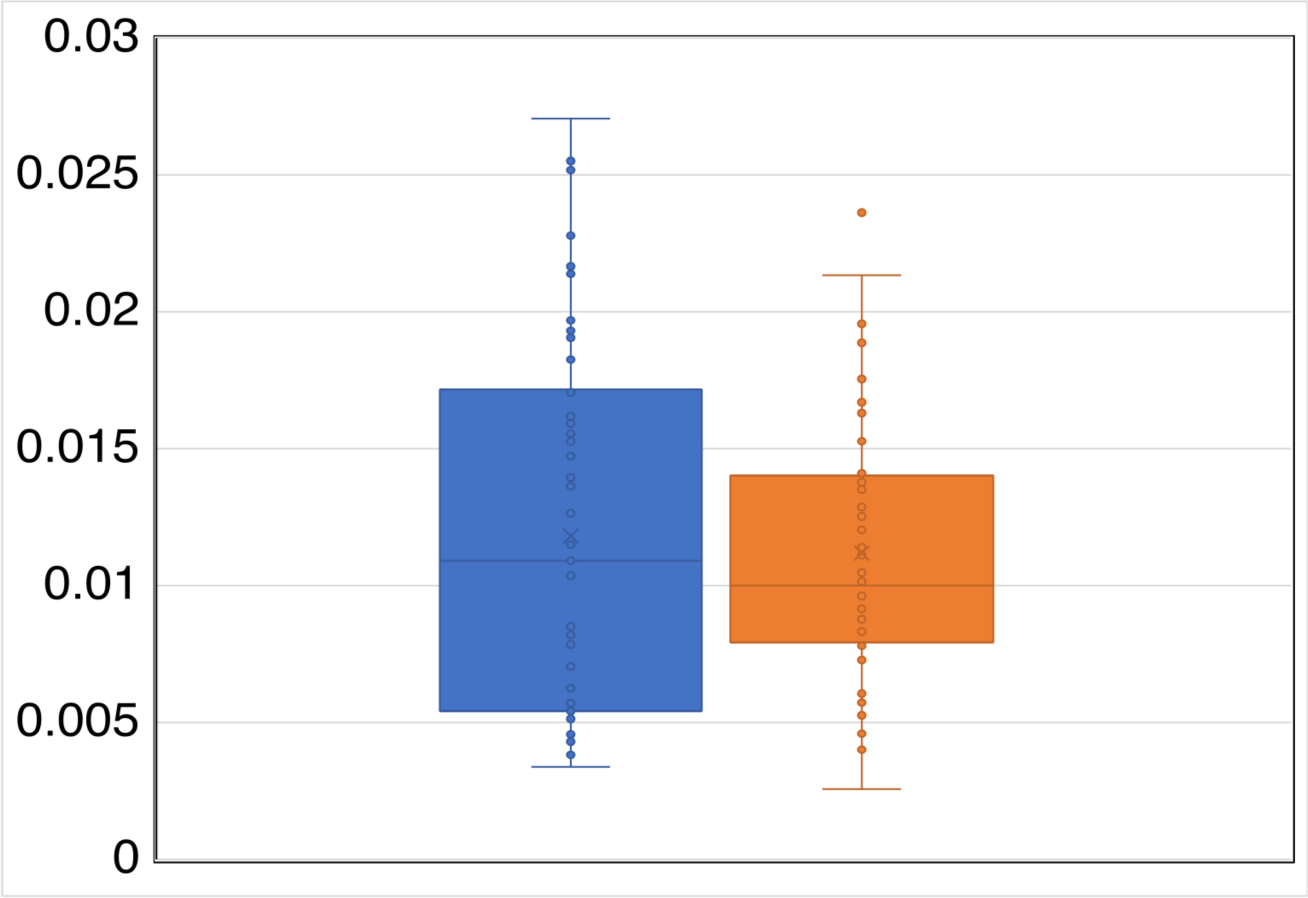
Proteins involved in extracellular activities, as a fraction of the total genes in the in the genome, for generalists (Blue) and specialists (Orange) taxa identified by this study. The difference is significant at p < 0.05 (Mann-Whitney test).

**Figure S14.**
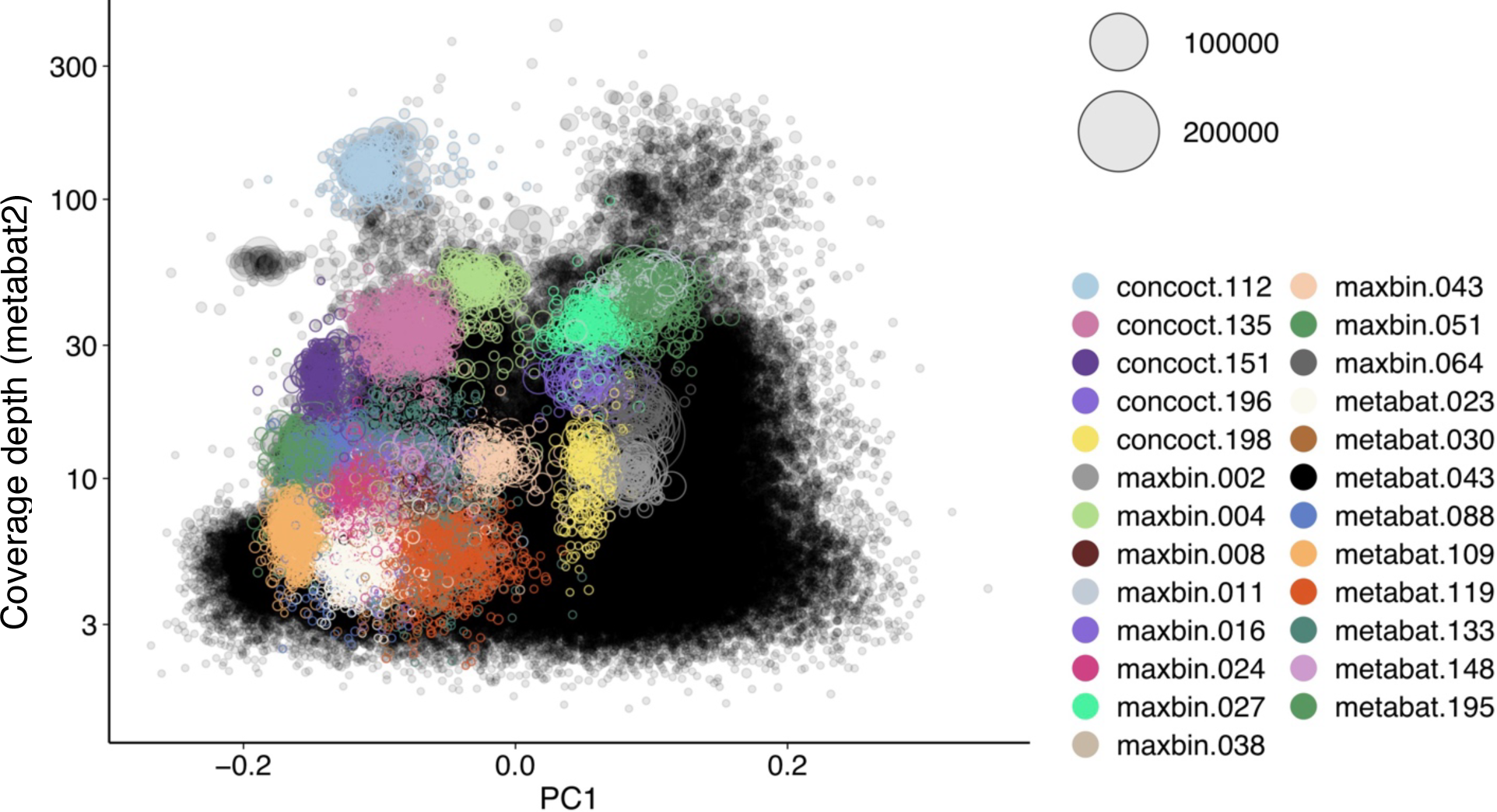
Contig coverage depth vs. principal component 1 of tetranucleotide frequencies for sample pico127 showed that each MAG we binned represents species-like, sequence clusters. Binned contigs by 3 different pieces of binning software (MaxBin2, MetaBAT2 and CONCOCT) are labelled by different color. The size of each circle represents the contig length.

**Figure S15.**
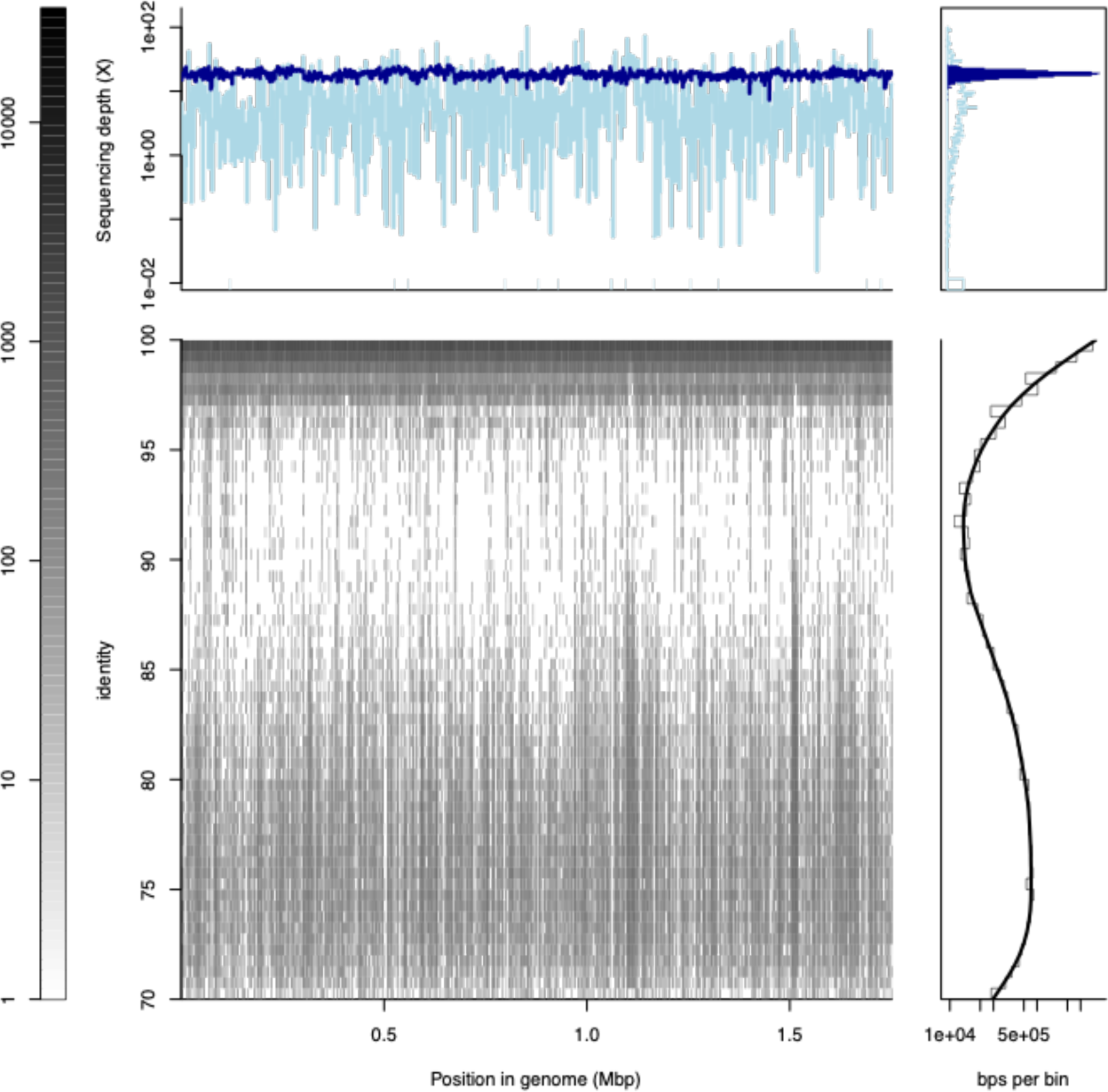
Recruitment plot of one MAG from sample pico497 (pico497_pico497.23.fasta) as an example that the binned MAGs represent sequence discrete populations. The reference MAG represents a sequence-discrete population in the pico 497 metagenome because there are many reads mapping on the MAG with >95% nucleotide, contrasting with reads showing 85-95% identity that are sparse. Average sequence coverage depth of this MAG is ∼23X. An interactive version (up left and bottom left panel) of this plot is available: https://github.com/jianshu93/RecruitmentPlot_blast/blob/main/example_out/pico497.23.html.zip (download and then open it in a browser). All Recruitment plots for all MAGs of each sample are available here: https://github.com/jianshu93/RecruitmentPlot_blast/tree/main/example_out/pico_rec_plot_2 and here: https://github.com/jianshu93/RecruitmentPlot_blast/tree/main/example_out/pico_rec_plot.

**Figure S16.**
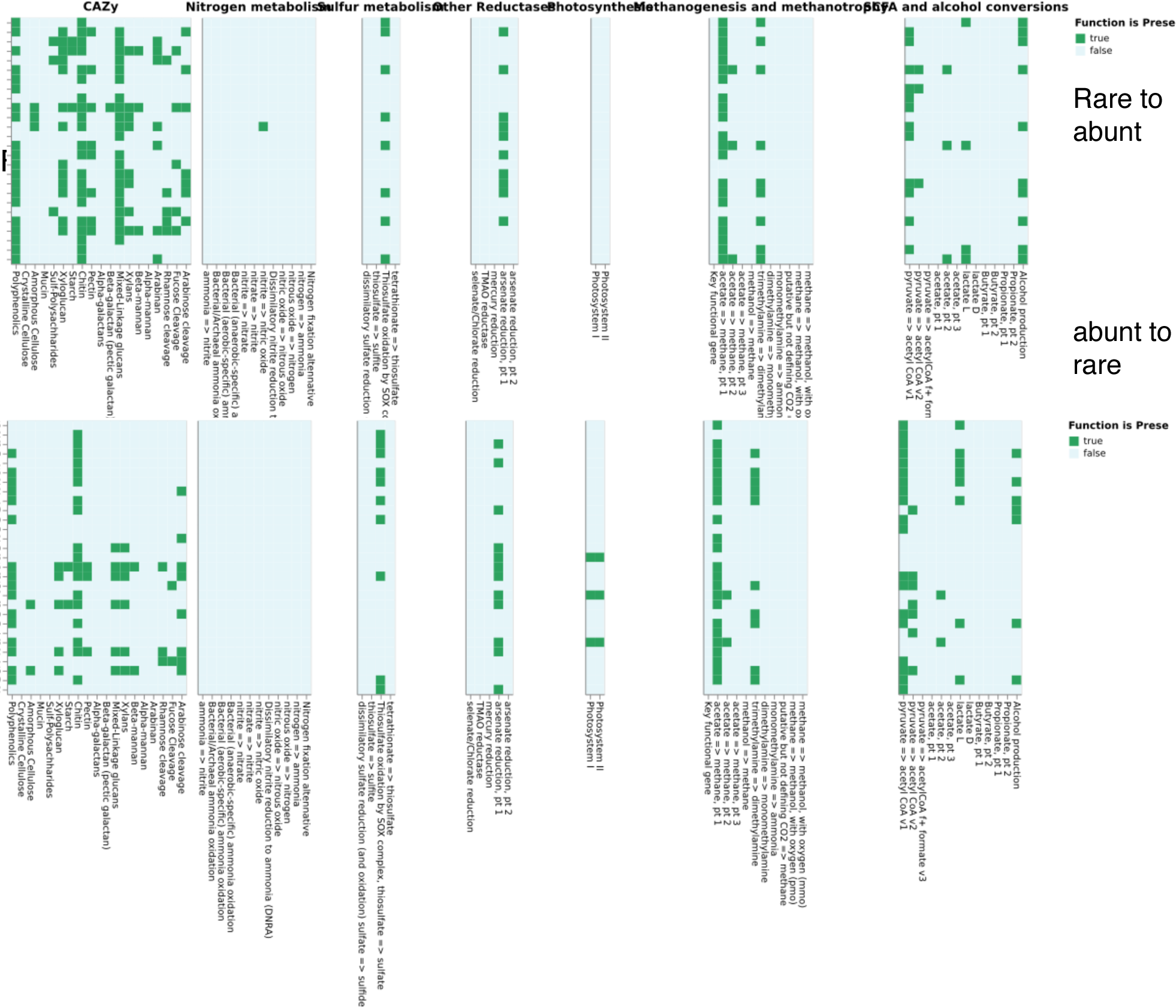
Metabolic pathways encoded in the genome of abundant-to-rare and rare-to-abundant MAGs for disturbance event 5 based on the DRAM software.

**Figure 17.**
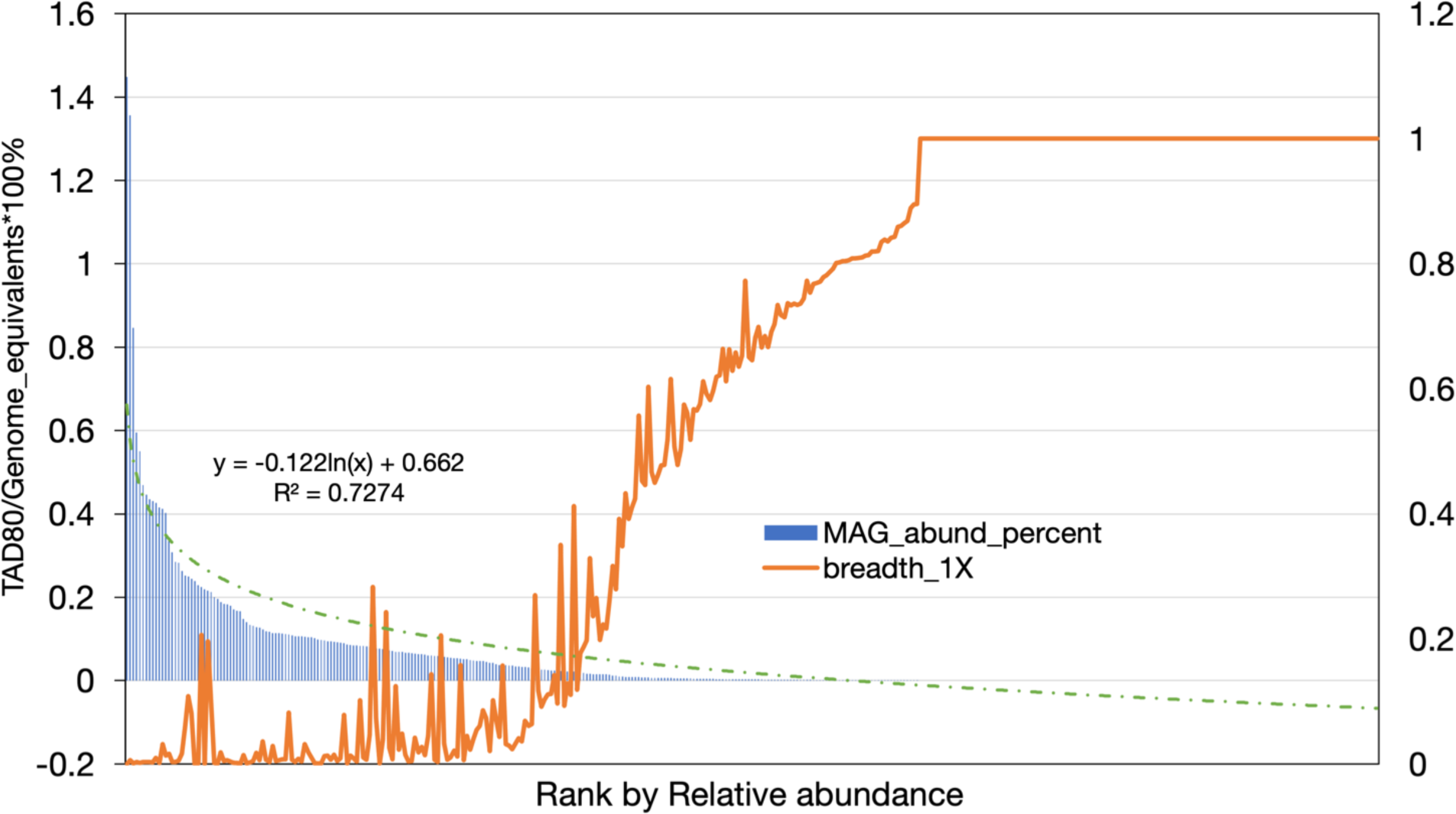
MAG sequence depth (left y axis, blue bar) and breadth (right y axis, orange line, shown as 1-coverage breadth) coverage distribution for pico127 before subsampling shows similar fitted line as the subsampled dataset and a slightly shifted abundance threshold for defining rare taxa (around 0.05).

**Figure S18.**
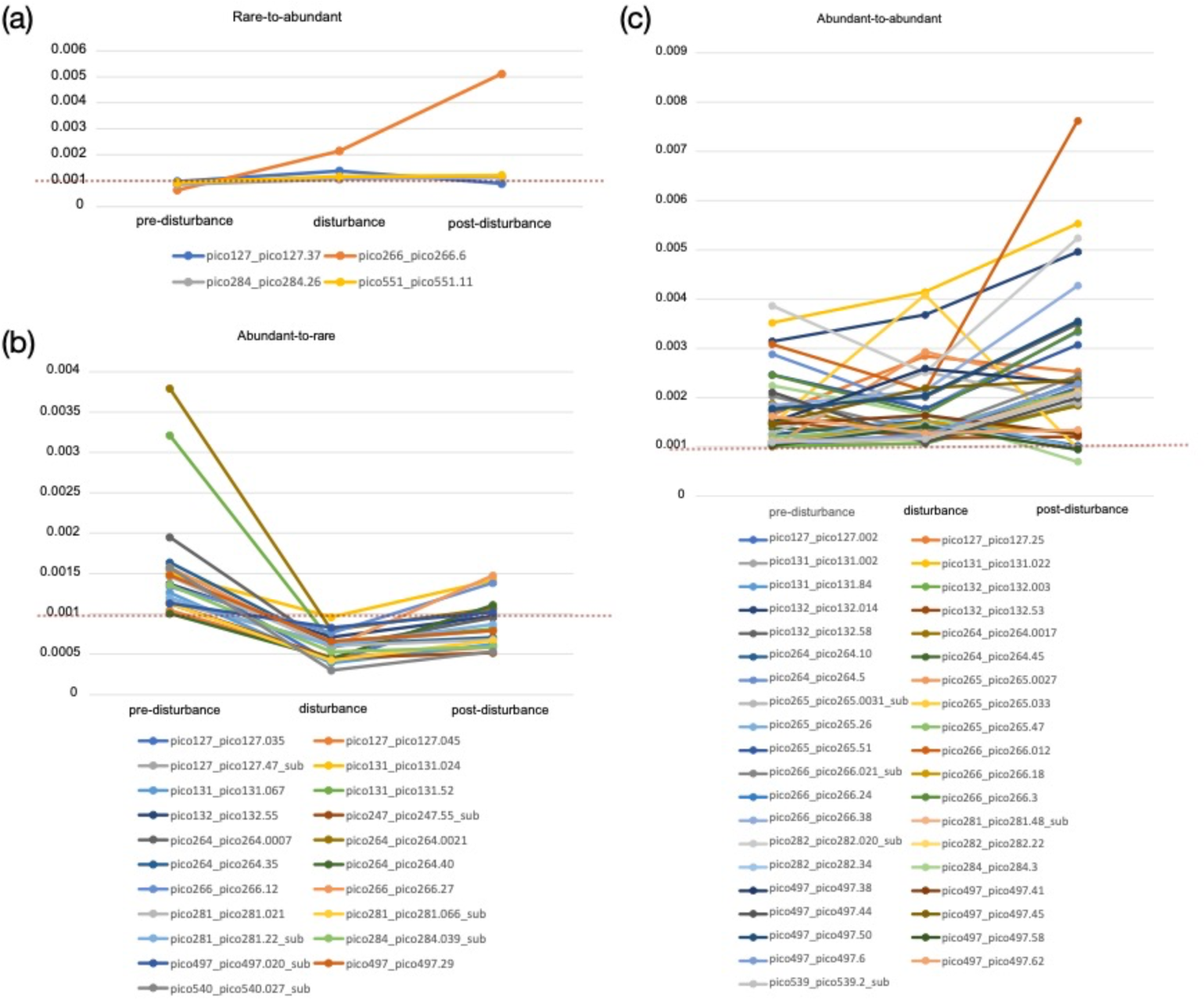
Relative abundance of MAGs assigned to rare to abundant (a), abundant to rare (b) and abundant to abundant (c) categories for disturbance event #3 (winter12_rainy_3 in Figure S1). Relative abundance (y-axis) was estimated as TAD80 sequence depth divided by genome equivalents to normalize for any average genome size differences between the samples as described in the Materials and Methods section. The red dashed line represents the threshold used to define rare taxa. Note the higher similarity in abundances between the pre- and post-disturbance samples relative to the disturbance sample, which indicates that stochastic processes have limited effect in identifying MAGs that change abundance categories due to the disturbance event.

**Figure S19.**
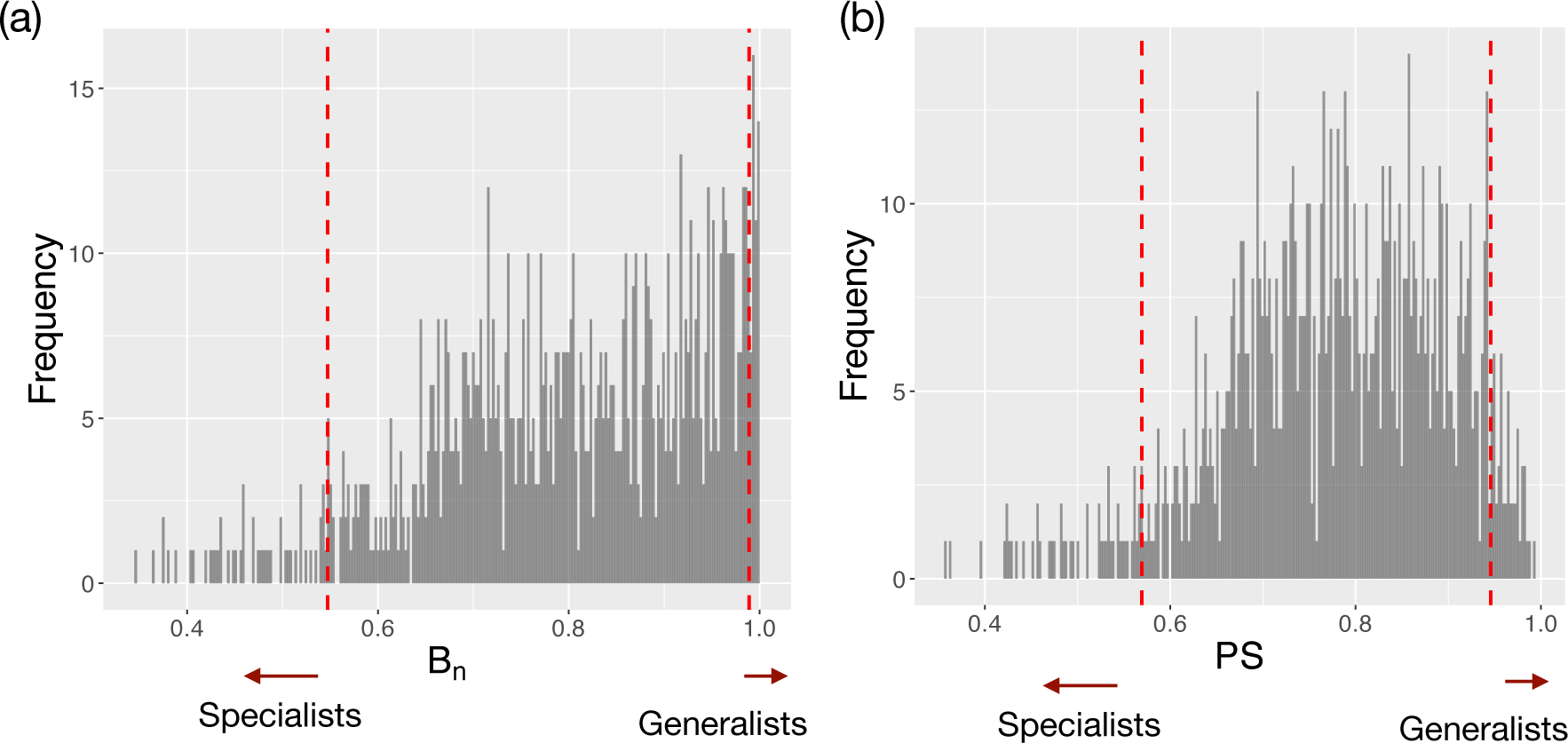
Levin’s Breadth index (a) and PS index (b) in MicroNiche package to identify generalists and specialists.

## Supplementary Tables

**Table S1.** Measured environmental variables for the time-series samples used in this study. Boldface numbers denote samples that associated with disturbance events. See Figure S1 for summary of each event.

|  | DayLength | Insolation | Temp<br>p | MLLW<sup>a</sup> | Barometric<br>Pressure | Salinity | Oxygen | OxygenS<br>aturation | pH | DIC | Chlor<br>o | NH4 | NOx<sup>b</sup> | PO4 | SiO4 |
| --- | --- | --- | --- | --- | --- | --- | --- | --- | --- | --- | --- | --- | --- | --- | --- |
| pico127 | 10.18 | 231.74 | 7.4 | 0.62 | 1005.3 | 32 | 9.74 | 99.8 | 7.92 | 1971 | 3.61 | 390.63 | 0.02 | 0.05 | 1.79 |
| **pico131** | **10.83** | **278.76** | **8.9** | **0.03** | **1031.8** | **32** | **9.89** | **104.9** | **7.92** | **1967.62** | **4.1** | **167.46** | **0** | **0.04** | **1.92** |
| pico132 | 11.07 | 296.61 | 9.9 | 0.89 | 1026.7 | 35 | 9.37 | 103.5 | 7.93 | 2073.19 | 2.45 | 174.05 | 0 | 0.04 | 0.92 |
| pico244 | 13.28 | 431.24 | 29 | 1.13 | 1020.9 | 34 | 6.55 | 103 | 8.08 | 2106.39 | 3.26 | 193.37 | 0 | 0.07 | 2.42 |
| **pico245** | **13.05** | **418.88** | **27.8** | **0.31** | **1019** | **35** | **5.85** | **90** | **8.02** | **2156.02** | **5.81** | **319.94** | **0** | **0.13** | **4.36** |
| pico247 | 12.61 | 392.56 | 28 | 0.25 | 1012.9 | 31 | 6.39 | 97.1 | 7.94 | 1986.3 | 7.96 | 218.94 | 0.09 | 0.03 | 13.95 |
| pico264 | 9.74 | 199.91 | 10 | 0.08 | 1018.7 | 31 | 8.8 | 94.3 | 7.92 | 2096.6 | 3.11 | 73.76 | 0.12 | 0 | 2.83 |
| **pico265** | **9.83** | **206.67** | **11.4** | **0.8** | **1019.6** | **33.5** | **8.73** | **98.1** | **7.92** | **2145.7** | **2.4** | **303.85** | **0.19** | **0** | **2.05** |
| pico266 | 9.96 | 215.95 | 9.6 | 0.2 | 1024.7 | 33 | 9.16 | 99.2 | 7.89 | 2157.37 | 2.38 | 229.76 | 0.1 | 0.01 | 3.58 |
| pico281 | 13.45 | 448.19 | 21.1 | -0.05 | 1023.5 | 30.5 | 7.74 | 103.1 | 7.87 | 2060.23 | 5.65 | 153 | 0.06 | 0.06 | 8.25 |
| **pico282** | **13.65** | **457.68** | **21.8** | **1.12** | **1017.2** | **34** | **7.84** | **106.8** | **7.95** | **2157.2** | **3.86** | **130** | **0.13** | **0** | **3.64** |
| pico284 | 13.99 | 472.05 | 22.3 | 0.94 | 1013.2 | 33 | 7.39 | 102.8 | 7.92 | 2119.8 | 5.9 | 123.5 | 0.03 | 0.02 | 7.52 |
| pico304 | 14.08 | 471.45 | 29.1 | 0.36 | 1016 | 33 | 6.74 | 107.2 | 8.01 | 2135.5 | 6.1 | 102 | 0 | 0 | 5.52 |
| pico497 | 10.52 | 256.02 | 10.1 | 0 | 1021.3 | 32.75 | 7.99 | 86.9 | 7.9 | 2115.77 | 1.82 | 24.5 | 0 | 0.02 | 1.94 |
| pico539 | 14.13 | 477.05 | 23 | 0.83 | 1026.1 | 34.78 | 7.03 | 97.7 | 7.94 | 2111.17 | 3.02 | 22 | 0.04 | 0.05 | 1.55 |
| **pico540** | **14.23** | **480.63** | **24.7** | **0.18** | **1020.3** | **34.57** | **6.65** | **95.1** | **7.95** | **2107.43** | **6.8** | **9** | **0.04** | **0.02** | **1.98** |
| pico550 | 13.57 | 446.17 | 26.9 | 0.84 | 1020 | 34.99 | 6.25 | 94.7 | 7.95 | 2099.73 | 5.52 | 29.5 | 0 | 0.03 | 4.52 |
| pico551 | 13.35 | 435.45 | 28.3 | 0.48 | 1013.8 | 35.41 | 5.73 | 89 | 7.97 | 2115.97 | 6.71 | 95 | 0.03 | 0.05 | 6.76 |
| **pico552** | **13.13** | **423.42** | **26.9** | **0.89** | **1024** | **34.97** | **6.26** | **94.7** | **7.95** | **2106.8** | **5.46** | **53** | **0** | **0.03** | **5.29** |

<sup>a</sup>mean of the two low tides.

**Table S2.**
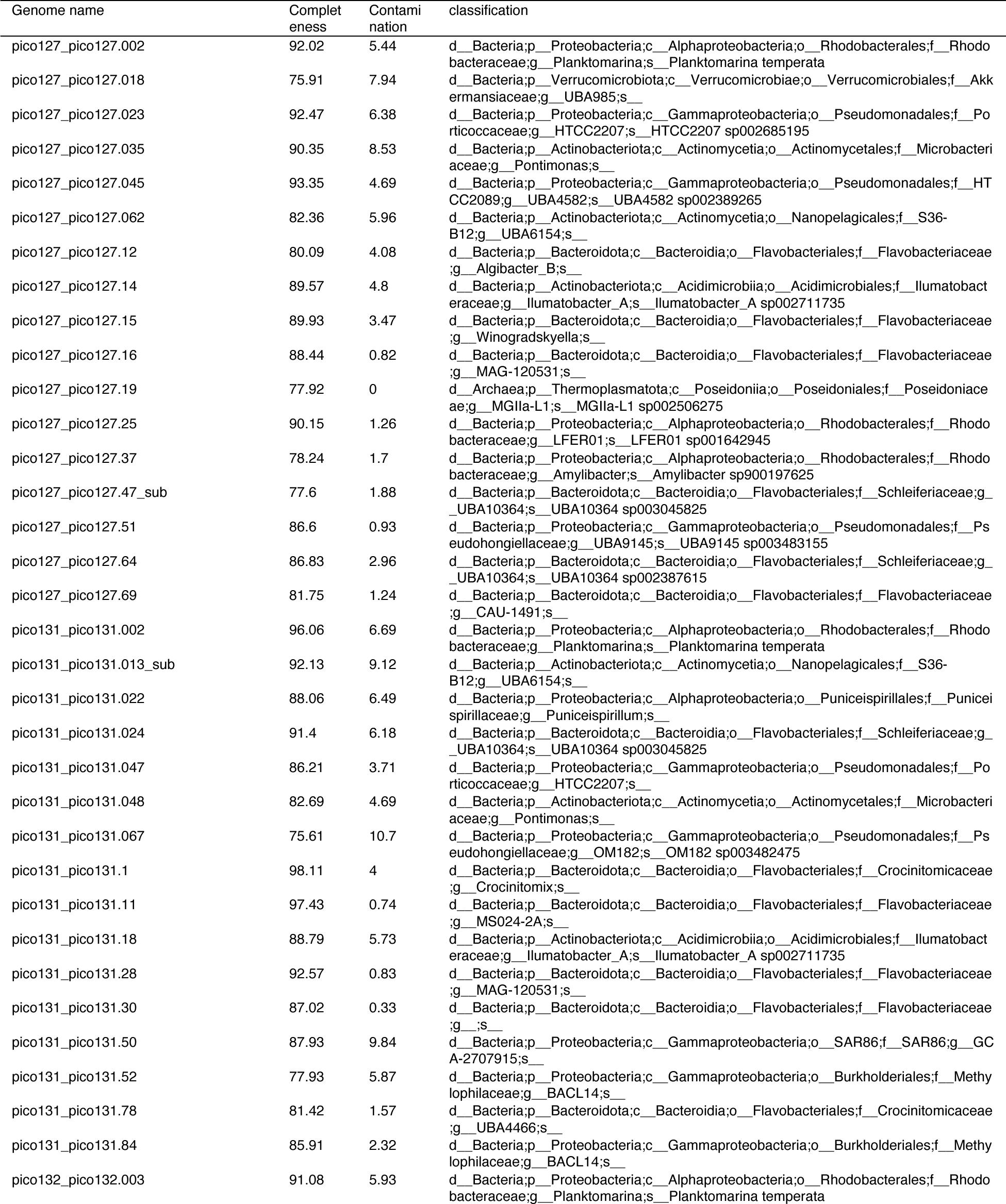
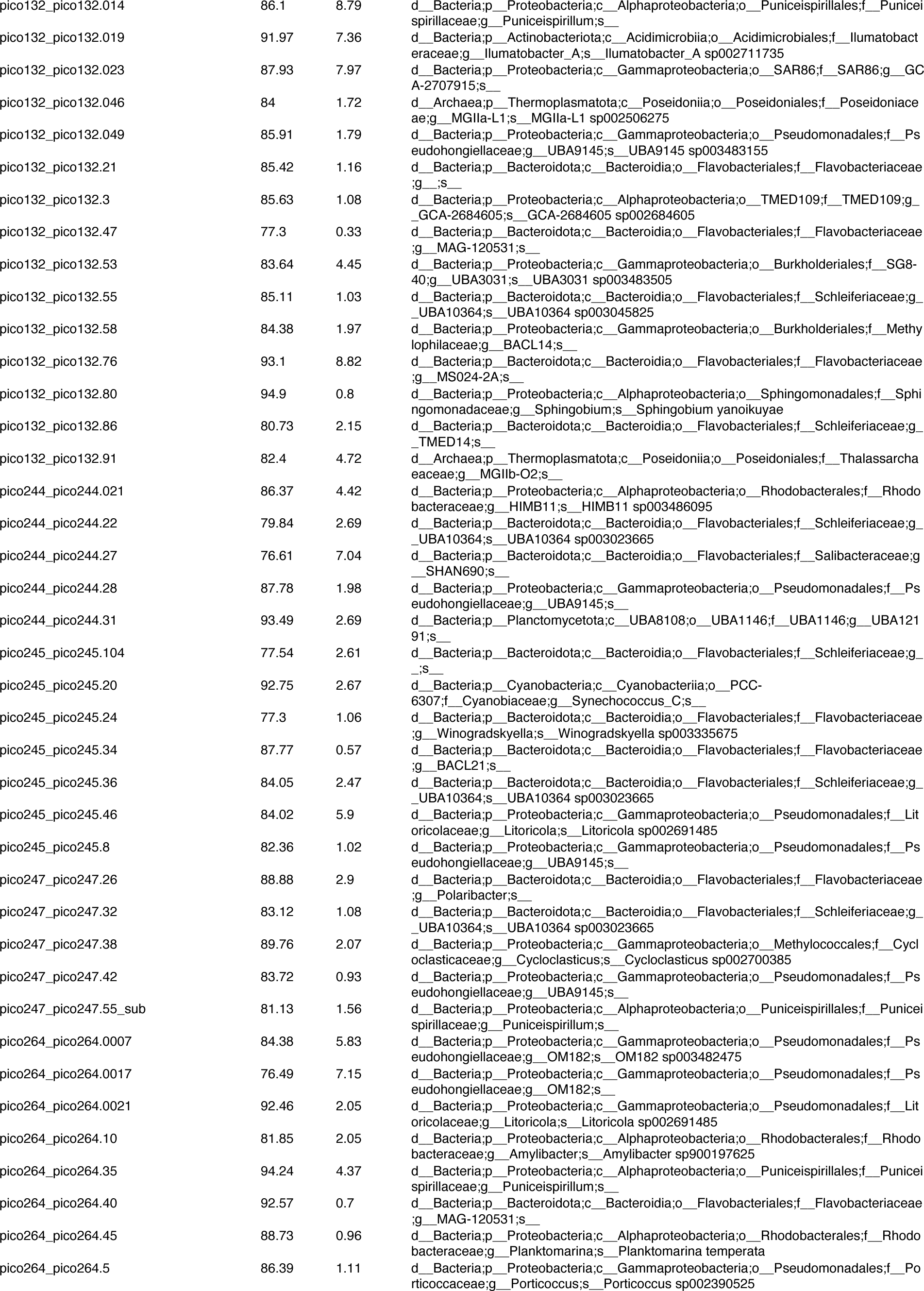
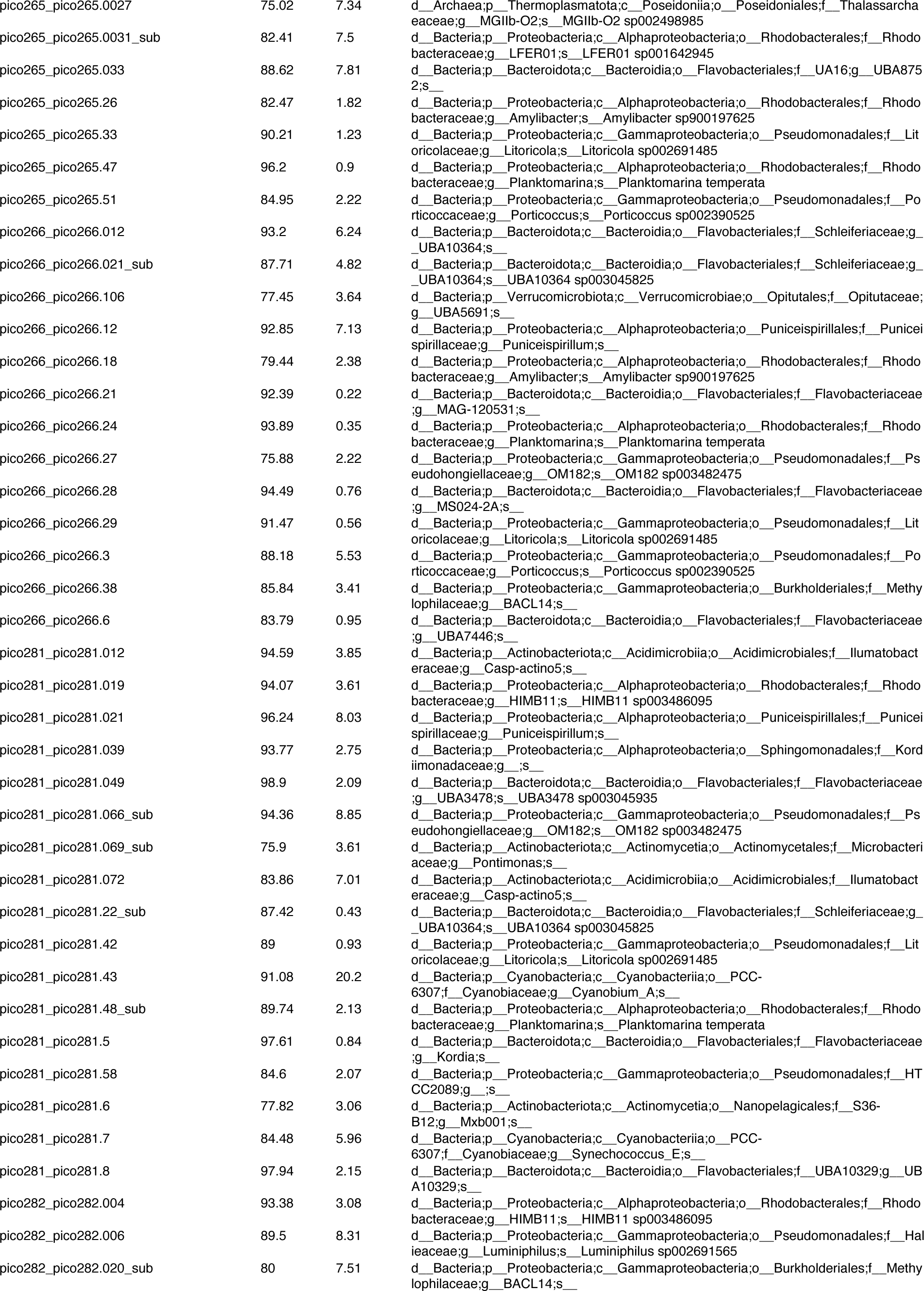
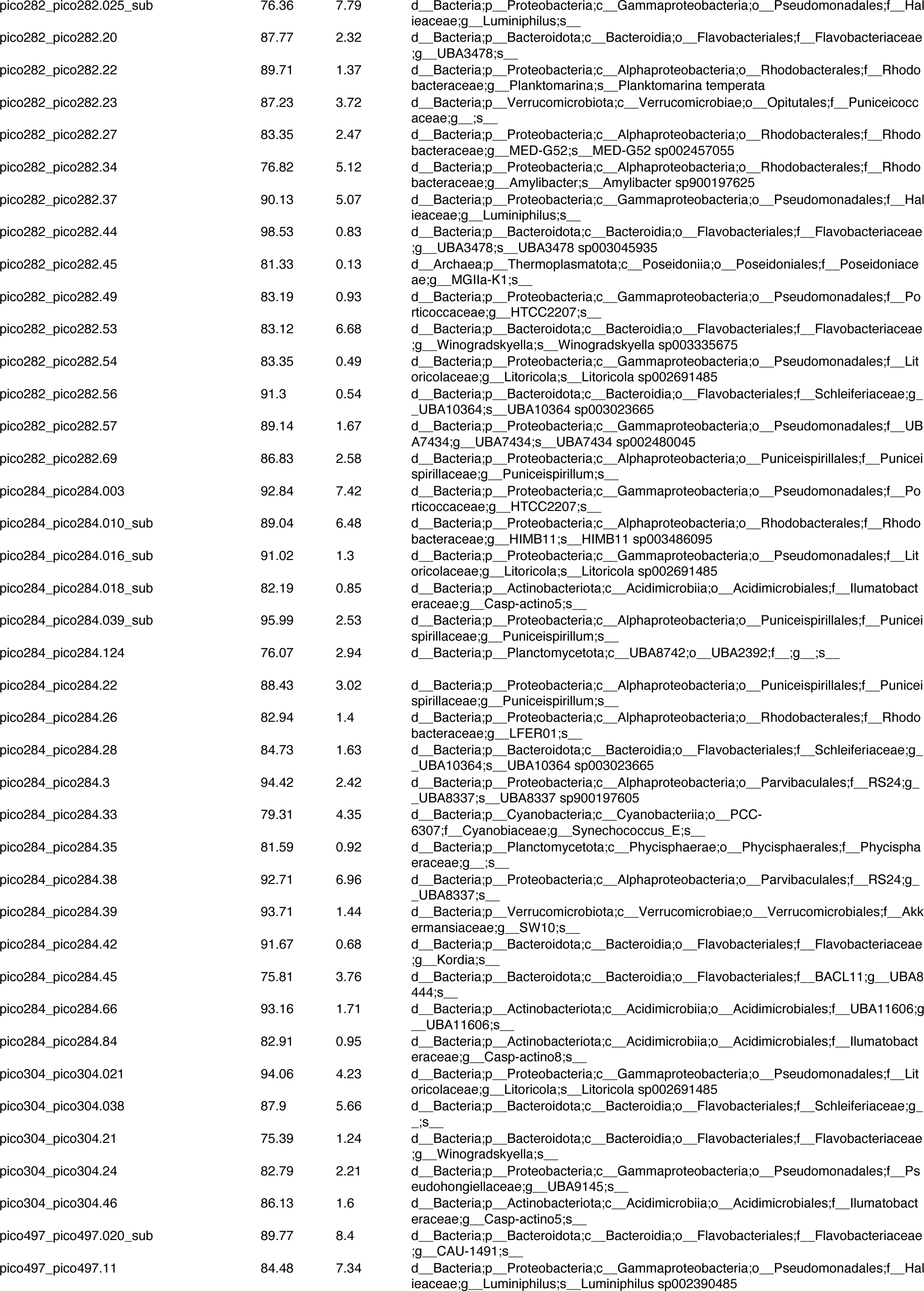
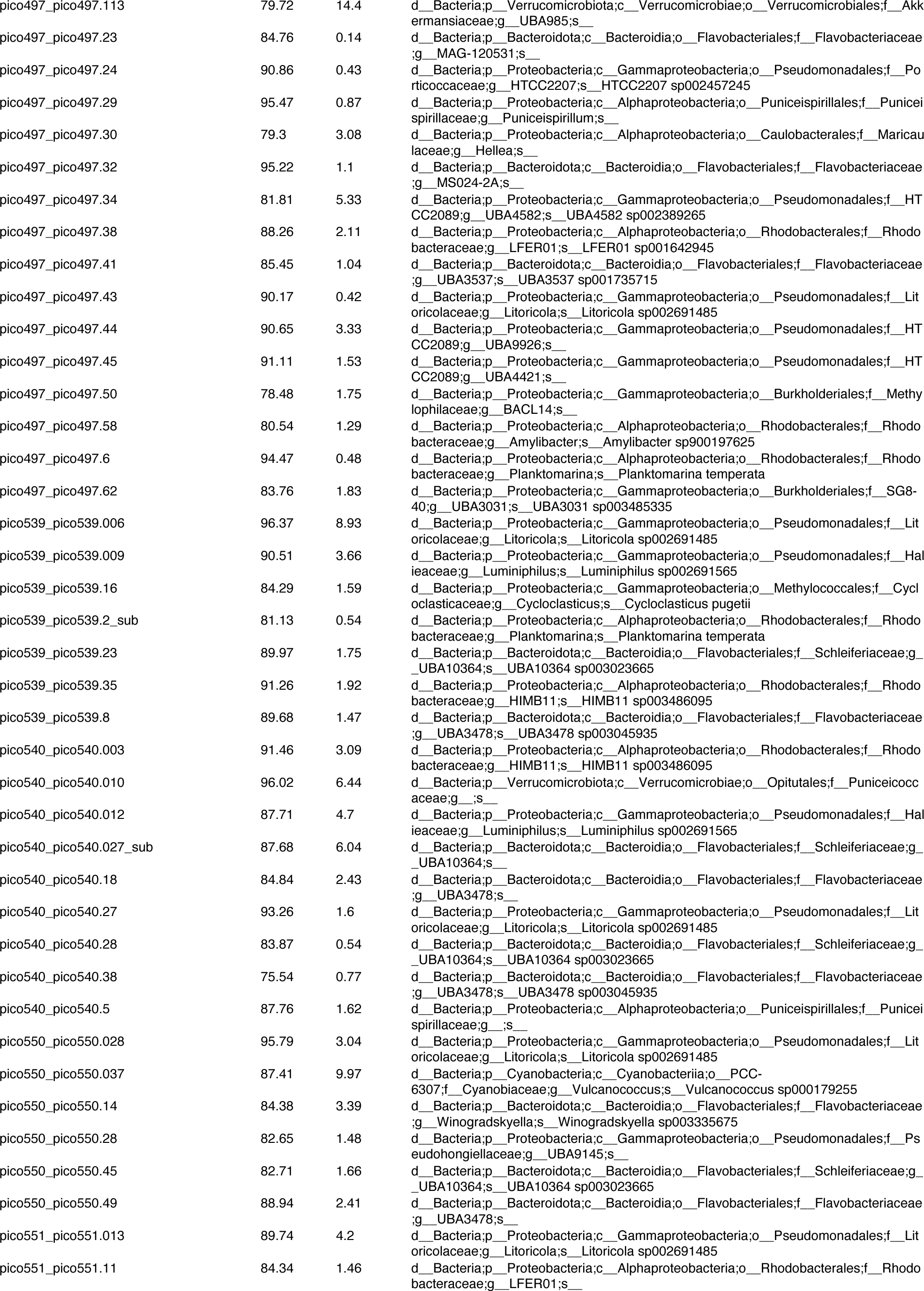
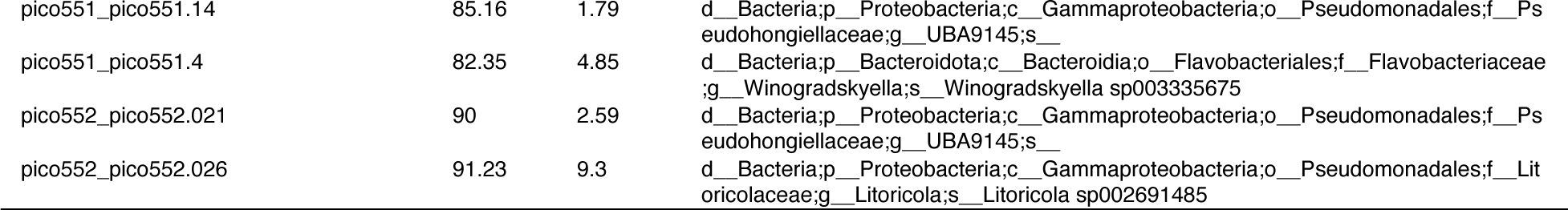
Quality and classification of the dereplicated, high quality MAGs used in the study. Quality was assessed based on CheckM results while classification is from GTDB-tk v1.1 against GTDB v207.

**Table S3.** Proportional Similarity index for all dereplicated, high quality MAGs. Main type and subtype of PS index are calculated according to the equation in the Material and Methods section.

|  | PS_maintype | Rank_transformed_PS_maintype | PS_subtype | Rank_transformed_PS_subtype | Rank_product | g_s |
| --- | --- | --- | --- | --- | --- | --- |
| pico127_pico127.002 | 0.423458459 | 56 | 0.42345846 | 69 | 3864 | specialists |
| pico127_pico127.018 | 0.388136546 | 36 | 0.26391317 | 26 | 936 | specialists |
| pico127_pico127.023 | 0.368573156 | 19 | 0.21413539 | 22 | 418 | specialists |
| pico127_pico127.035 | 0.63576366 | 126 | 0.52752312 | 115 | 14490 | potential_specialists |
| pico127_pico127.045 | 0.368421053 | 10 | 0.36842105 | 47 | 470 | specialists |
| pico127_pico127.062 | 0.431132576 | 64 | 0.3110947 | 31 | 1984 | specialists |
| pico127_pico127.12 | 0.368421053 | 11 | 0.15789474 | 2 | 22 | specialists |
| pico127_pico127.14 | 0.374401623 | 30 | 0.19480945 | 15 | 450 | specialists |
| pico127_pico127.15 | 0.396249494 | 41 | 0.18572318 | 13 | 533 | specialists |
| pico127_pico127.16 | 0.452641622 | 74 | 0.45264162 | 86 | 6364 | potential_specialists |
| pico127_pico127.19 | 0.369623126 | 21 | 0.2138458 | 21 | 441 | specialists |
| pico127_pico127.25 | 0.562866455 | 106 | 0.51621983 | 109 | 11554 | potential_specialists |
| pico127_pico127.37 | 0.425467381 | 58 | 0.42546738 | 71 | 4118 | potential_specialists |
| pico127_pico127.47_sub | 0.422113545 | 55 | 0.36605702 | 46 | 2530 | specialists |
| pico127_pico127.51 | 0.370762565 | 26 | 0.35848922 | 45 | 1170 | specialists |
| pico127_pico127.64 | 0.379665689 | 32 | 0.22604431 | 25 | 800 | specialists |
| pico127_pico127.69 | 0.473121423 | 83 | 0.33280089 | 39 | 3237 | specialists |
| pico131_pico131.002 | 0.436110027 | 65 | 0.43611003 | 77 | 5005 | potential_specialists |
| pico131_pico131.013_sub | 0.420230952 | 53 | 0.31395254 | 32 | 1696 | specialists |
| pico131_pico131.022 | 0.561159425 | 104 | 0.56115943 | 123 | 12792 | potential_specialists |
| pico131_pico131.024 | 0.507118015 | 94 | 0.50711802 | 108 | 10152 | potential_specialists |
| pico131_pico131.047 | 0.379613634 | 31 | 0.27880432 | 28 | 868 | specialists |
| pico131_pico131.048 | 0.569082781 | 108 | 0.43072985 | 76 | 8208 | potential_specialists |
| pico131_pico131.067 | 0.53950108 | 99 | 0.53950108 | 117 | 11583 | potential_specialists |
| pico131_pico131.1 | 0.368421053 | 12 | 0.15789474 | 3 | 36 | specialists |
| pico131_pico131.11 | 0.399710489 | 44 | 0.39971049 | 60 | 2640 | specialists |
| pico131_pico131.18 | 0.374148902 | 29 | 0.19081107 | 14 | 406 | specialists |
| pico131_pico131.28 | 0.477135272 | 87 | 0.47713527 | 103 | 8961 | potential_specialists |
| pico131_pico131.30 | 0.369948435 | 23 | 0.2088105 | 19 | 437 | specialists |
| pico131_pico131.50 | 0.369439545 | 20 | 0.17008161 | 9 | 180 | specialists |
| pico131_pico131.52 | 0.650916648 | 135 | 0.58219334 | 128 | 17280 | potential_specialists |
| pico131_pico131.78 | 0.501790492 | 91 | 0.42672819 | 73 | 6643 | potential_specialists |
| pico131_pico131.84 | 0.400183247 | 45 | 0.40018325 | 61 | 2745 | specialists |
| pico132_pico132.003 | 0.430605081 | 62 | 0.43060508 | 74 | 4588 | potential_specialists |
| pico132_pico132.014 | 0.557862425 | 102 | 0.55786243 | 121 | 12342 | potential_specialists |
| pico132_pico132.019 | 0.370491173 | 24 | 0.17545465 | 12 | 288 | specialists |
| pico132_pico132.023 | 0.37049483 | 25 | 0.17157076 | 10 | 250 | specialists |
| pico132_pico132.046 | 0.373272922 | 28 | 0.21114827 | 20 | 560 | specialists |
| pico132_pico132.049 | 0.372330492 | 27 | 0.35660097 | 44 | 1188 | specialists |
| pico132_pico132.21 | 0.369896297 | 22 | 0.19913859 | 16 | 352 | specialists |
| pico132_pico132.47 | 0.475200469 | 85 | 0.47520047 | 100 | 8500 | potential_specialists |
| pico132_pico132.53 | 0.631578947 | 120 | 0.57501467 | 126 | 15120 | potential_specialists |
| pico132_pico132.55 | 0.531840462 | 97 | 0.53184046 | 116 | 11252 | potential_specialists |
| pico132_pico132.58 | 0.400614931 | 46 | 0.40061493 | 62 | 2852 | specialists |
| pico132_pico132.76 | 0.410929366 | 50 | 0.41092937 | 65 | 3250 | specialists |
| pico132_pico132.80 | 0.368421053 | 13 | 0.15789474 | 4 | 52 | specialists |
| pico132_pico132.86 | 0.368421053 | 14 | 0.15789474 | 5 | 70 | specialists |
| pico132_pico132.91 | 0.368421053 | 15 | 0.15789474 | 6 | 90 | specialists |
| pico244_pico244.021 | 0.583159757 | 111 | 0.58315976 | 130 | 14430 | potential_specialists |
| pico244_pico244.22 | 0.877163298 | 190 | 0.81938119 | 192 | 36480 | generalists |
| pico244_pico244.27 | 0.822616536 | 180 | 0.54485225 | 119 | 21420 | generalists |
| pico244_pico244.28 | 0.736842105 | 151 | 0.52308377 | 112 | 16912 | potential_specialists |
| pico244_pico244.31 | 0.368421053 | 16 | 0.31578947 | 33 | 528 | specialists |
| pico245_pico245.104 | 0.729576935 | 150 | 0.72957694 | 169 | 25350 | generalists |
| pico245_pico245.20 | 0.399304254 | 43 | 0.39930425 | 59 | 2537 | specialists |
| pico245_pico245.24 | 0.848327247 | 184 | 0.80004032 | 184 | 33856 | generalists |
| pico245_pico245.34 | 0.411401648 | 51 | 0.41140165 | 66 | 3366 | specialists |
| pico245_pico245.36 | 0.785388171 | 164 | 0.73923476 | 171 | 28044 | generalists |
| pico245_pico245.46 | 0.873322775 | 189 | 0.87332278 | 196 | 37044 | generalists |
| pico245_pico245.8 | 0.368421053 | 17 | 0.36842105 | 48 | 816 | specialists |
| pico247_pico247.26 | 0.430238065 | 61 | 0.32984125 | 38 | 2318 | specialists |
| pico247_pico247.32 | 0.786770663 | 165 | 0.74447828 | 172 | 28380 | generalists |
| pico247_pico247.38 | 0.708718616 | 148 | 0.60702685 | 138 | 20424 | generalists |
| pico247_pico247.42 | 0.736842105 | 152 | 0.5169276 | 111 | 16872 | potential_specialists |
| pico247_pico247.55_sub | 0.859885256 | 186 | 0.64950016 | 148 | 27528 | generalists |
| pico264_pico264.0007 | 0.583828953 | 112 | 0.58382895 | 132 | 14784 | potential_specialists |
| pico264_pico264.0017 | 0.631578947 | 116 | 0.54933814 | 120 | 13920 | potential_specialists |
| pico264_pico264.0021 | 0.742954057 | 156 | 0.69973034 | 163 | 25428 | generalists |
| pico264_pico264.10 | 0.424617986 | 57 | 0.42461799 | 70 | 3990 | specialists |
| pico264_pico264.35 | 0.690546661 | 147 | 0.65778919 | 151 | 22197 | generalists |
| pico264_pico264.40 | 0.453082912 | 75 | 0.45308291 | 87 | 6525 | potential_specialists |
| pico264_pico264.45 | 0.444382305 | 70 | 0.44438231 | 81 | 5670 | potential_specialists |
| pico264_pico264.5 | 0.390515182 | 38 | 0.39051518 | 54 | 2052 | specialists |
| pico265_pico265.0027 | 0.38236677 | 33 | 0.33500781 | 40 | 1320 | specialists |
| pico265_pico265.0031_sub | 0.552413487 | 101 | 0.45518174 | 90 | 9090 | potential_specialists |
| pico265_pico265.033 | 0.85593742 | 185 | 0.71532754 | 166 | 30710 | generalists |
| pico265_pico265.26 | 0.446349242 | 71 | 0.44634924 | 82 | 5822 | potential_specialists |
| pico265_pico265.33 | 0.80560775 | 175 | 0.80560775 | 186 | 32550 | generalists |
| pico265_pico265.47 | 0.443322504 | 69 | 0.4433225 | 80 | 5520 | potential_specialists |
| pico265_pico265.51 | 0.39330795 | 39 | 0.39330795 | 55 | 2145 | specialists |
| pico266_pico266.012 | 0.497613626 | 90 | 0.39715618 | 58 | 5220 | potential_specialists |
| pico266_pico266.021_sub | 0.51642883 | 95 | 0.51642883 | 110 | 10450 | potential_specialists |
| pico266_pico266.106 | 0.477251097 | 88 | 0.46489989 | 95 | 8360 | potential_specialists |
| pico266_pico266.12 | 0.744500514 | 157 | 0.58240823 | 129 | 20253 | generalists |
| pico266_pico266.18 | 0.446894707 | 72 | 0.44689471 | 83 | 5976 | potential_specialists |
| pico266_pico266.21 | 0.467877787 | 81 | 0.46787779 | 96 | 7776 | potential_specialists |
| pico266_pico266.24 | 0.43068744 | 63 | 0.43068744 | 75 | 4725 | potential_specialists |
| pico266_pico266.27 | 0.560495195 | 103 | 0.5604952 | 122 | 12566 | potential_specialists |
| pico266_pico266.28 | 0.416246309 | 52 | 0.41624631 | 67 | 3484 | specialists |
| pico266_pico266.29 | 0.790423224 | 168 | 0.79042322 | 180 | 30240 | generalists |
| pico266_pico266.3 | 0.395619079 | 40 | 0.39561908 | 57 | 2280 | specialists |
| pico266_pico266.38 | 0.403594818 | 49 | 0.40359482 | 64 | 3136 | specialists |
| pico266_pico266.6 | 0.421489444 | 54 | 0.3944986 | 56 | 3024 | specialists |
| pico281_pico281.012 | 0.562446661 | 105 | 0.45012795 | 84 | 8820 | potential_specialists |
| pico281_pico281.019 | 0.681197855 | 146 | 0.68119786 | 160 | 23360 | generalists |
| pico281_pico281.021 | 0.784992584 | 163 | 0.58052541 | 127 | 20701 | generalists |
| pico281_pico281.039 | 0.263157895 | 1 | 0.15789474 | 7 | 7 | specialists |
| pico281_pico281.049 | 0.649566718 | 134 | 0.63816212 | 145 | 19430 | generalists |
| pico281_pico281.066_sub | 0.637376654 | 127 | 0.58376149 | 131 | 16637 | potential_specialists |
| pico281_pico281.069_sub | 0.661892478 | 139 | 0.62372096 | 142 | 19738 | generalists |
| pico281_pico281.072 | 0.803858908 | 173 | 0.67049961 | 155 | 26815 | generalists |
| pico281_pico281.22_sub | 0.631578947 | 117 | 0.56390397 | 124 | 14508 | potential_specialists |
| pico281_pico281.42 | 0.796773633 | 171 | 0.79677363 | 183 | 31293 | generalists |
| pico281_pico281.43 | 0.451883531 | 73 | 0.35607904 | 43 | 3139 | specialists |
| pico281_pico281.48_sub | 0.453392708 | 76 | 0.45339271 | 88 | 6688 | potential_specialists |
| pico281_pico281.5 | 0.263157895 | 2 | 0.15789474 | 8 | 16 | specialists |
| pico281_pico281.58 | 0.456705756 | 78 | 0.45670576 | 92 | 7176 | potential_specialists |
| pico281_pico281.6 | 0.551484855 | 100 | 0.37016243 | 50 | 5000 | potential_specialists |
| pico281_pico281.7 | 0.640098372 | 130 | 0.58441921 | 134 | 17420 | potential_specialists |
| pico281_pico281.8 | 0.631578947 | 118 | 0.31578947 | 34 | 4012 | potential_specialists |
| pico282_pico282.004 | 0.6403223 | 131 | 0.6403223 | 147 | 19257 | generalists |
| pico282_pico282.006 | 0.671593237 | 142 | 0.67159324 | 156 | 22152 | generalists |
| pico282_pico282.020_sub | 0.828916846 | 182 | 0.8168228 | 189 | 34398 | generalists |
| pico282_pico282.025_sub | 0.663334117 | 140 | 0.6325504 | 143 | 20020 | generalists |
| pico282_pico282.20 | 0.664713248 | 141 | 0.66471325 | 153 | 21573 | generalists |
| pico282_pico282.22 | 0.45686551 | 79 | 0.45686551 | 93 | 7347 | potential_specialists |
| pico282_pico282.23 | 0.291370205 | 6 | 0.29137021 | 29 | 174 | specialists |
| pico282_pico282.27 | 0.677997507 | 145 | 0.67799751 | 159 | 23055 | generalists |
| pico282_pico282.34 | 0.493896425 | 89 | 0.49389643 | 106 | 9434 | potential_specialists |
| pico282_pico282.37 | 0.635662941 | 125 | 0.45667272 | 91 | 11375 | potential_specialists |
| pico282_pico282.44 | 0.814183258 | 177 | 0.69188627 | 162 | 28674 | generalists |
| pico282_pico282.45 | 0.632700831 | 122 | 0.5434132 | 118 | 14396 | potential_specialists |
| pico282_pico282.49 | 0.278869412 | 4 | 0.22598072 | 24 | 96 | specialists |
| pico282_pico282.53 | 0.871388376 | 188 | 0.81458879 | 188 | 35344 | generalists |
| pico282_pico282.54 | 0.777679987 | 161 | 0.77461966 | 176 | 28336 | generalists |
| pico282_pico282.56 | 0.909869328 | 196 | 0.84599718 | 194 | 38024 | generalists |
| pico282_pico282.57 | 0.573632863 | 110 | 0.37437544 | 51 | 5610 | potential_specialists |
| pico282_pico282.69 | 0.585079555 | 113 | 0.58507956 | 135 | 15255 | potential_specialists |
| pico284_pico284.003 | 0.271301532 | 3 | 0.22113624 | 23 | 69 | specialists |
| pico284_pico284.010_sub | 0.652653101 | 137 | 0.6526531 | 150 | 20550 | generalists |
| pico284_pico284.016_sub | 0.809456719 | 176 | 0.73818279 | 170 | 29920 | generalists |
| pico284_pico284.018_sub | 0.564506002 | 107 | 0.45206288 | 85 | 9095 | potential_specialists |
| pico284_pico284.039_sub | 0.794761335 | 170 | 0.61182428 | 139 | 23630 | generalists |
| pico284_pico284.124 | 0.537085871 | 98 | 0.31578947 | 35 | 3430 | specialists |
| pico284_pico284.22 | 0.59071919 | 114 | 0.59071919 | 137 | 15618 | potential_specialists |
| pico284_pico284.26 | 0.76364092 | 160 | 0.70299222 | 164 | 26240 | generalists |
| pico284_pico284.28 | 0.786840161 | 166 | 0.74531885 | 173 | 28718 | generalists |
| pico284_pico284.3 | 0.503421902 | 93 | 0.49997918 | 107 | 9951 | potential_specialists |
| pico284_pico284.33 | 0.641764445 | 132 | 0.58603094 | 136 | 17952 | generalists |
| pico284_pico284.35 | 0.279677509 | 5 | 0.17441435 | 11 | 55 | specialists |
| pico284_pico284.38 | 0.633017633 | 123 | 0.48839066 | 105 | 12915 | potential_specialists |
| pico284_pico284.39 | 0.296261077 | 8 | 0.27554128 | 27 | 216 | specialists |
| pico284_pico284.42 | 0.403237172 | 48 | 0.29797401 | 30 | 1440 | specialists |
| pico284_pico284.45 | 0.76130285 | 159 | 0.61406967 | 140 | 22260 | generalists |
| pico284_pico284.66 | 0.29618981 | 7 | 0.20305605 | 17 | 119 | specialists |
| pico284_pico284.84 | 0.45740534 | 80 | 0.35214218 | 42 | 3360 | specialists |
| pico304_pico304.021 | 0.882871728 | 191 | 0.78105113 | 178 | 33998 | generalists |
| pico304_pico304.038 | 0.82235419 | 179 | 0.81857435 | 191 | 34189 | generalists |
| pico304_pico304.21 | 0.903779827 | 194 | 0.88643037 | 197 | 38218 | generalists |
| pico304_pico304.24 | 0.736842105 | 154 | 0.52631579 | 114 | 17556 | generalists |
| pico304_pico304.46 | 0.634059936 | 124 | 0.62078295 | 141 | 17484 | generalists |
| pico497_pico497.020_sub | 0.468014967 | 82 | 0.46801497 | 97 | 7954 | potential_specialists |
| pico497_pico497.11 | 0.398377457 | 42 | 0.20406253 | 18 | 756 | specialists |
| pico497_pico497.113 | 0.790461853 | 169 | 0.75942498 | 174 | 29406 | generalists |
| pico497_pico497.23 | 0.475211368 | 86 | 0.47521137 | 101 | 8686 | potential_specialists |
| pico497_pico497.24 | 0.440497975 | 67 | 0.14141051 | 1 | 67 | specialists |
| pico497_pico497.29 | 0.777987443 | 162 | 0.68858344 | 161 | 26082 | generalists |
| pico497_pico497.30 | 0.425662299 | 59 | 0.32651485 | 37 | 2183 | specialists |
| pico497_pico497.32 | 0.390184283 | 37 | 0.34259029 | 41 | 1517 | specialists |
| pico497_pico497.34 | 0.368421053 | 18 | 0.36842105 | 49 | 882 | specialists |
| pico497_pico497.38 | 0.571407805 | 109 | 0.47611046 | 102 | 11118 | potential_specialists |
| pico497_pico497.41 | 0.627042251 | 115 | 0.58438548 | 133 | 15295 | potential_specialists |
| pico497_pico497.43 | 0.825211701 | 181 | 0.8252117 | 193 | 34933 | generalists |
| pico497_pico497.44 | 0.639132273 | 128 | 0.47278698 | 98 | 12544 | potential_specialists |
| pico497_pico497.45 | 0.517694348 | 96 | 0.46370467 | 94 | 9024 | potential_specialists |
| pico497_pico497.50 | 0.401854744 | 47 | 0.40185474 | 63 | 2961 | specialists |
| pico497_pico497.58 | 0.425943775 | 60 | 0.42594378 | 72 | 4320 | potential_specialists |
| pico497_pico497.6 | 0.436242542 | 66 | 0.43624254 | 78 | 5148 | potential_specialists |
| pico497_pico497.62 | 0.631710406 | 121 | 0.57427787 | 125 | 15125 | potential_specialists |
| pico539_pico539.006 | 0.454219914 | 77 | 0.45421991 | 89 | 6853 | potential_specialists |
| pico539_pico539.009 | 0.675638785 | 144 | 0.67563879 | 158 | 22752 | generalists |
| pico539_pico539.16 | 0.631578947 | 119 | 0.48242034 | 104 | 12376 | potential_specialists |
| pico539_pico539.2_sub | 0.440732288 | 68 | 0.44073229 | 79 | 5372 | potential_specialists |
| pico539_pico539.23 | 0.904921777 | 195 | 0.85020243 | 195 | 38025 | generalists |
| pico539_pico539.35 | 0.658872813 | 138 | 0.65887281 | 152 | 20976 | generalists |
| pico539_pico539.8 | 0.64862534 | 133 | 0.63402808 | 144 | 19152 | generalists |
| pico540_pico540.003 | 0.639564838 | 129 | 0.63956484 | 146 | 18834 | generalists |
| pico540_pico540.010 | 0.316447668 | 9 | 0.31644767 | 36 | 324 | specialists |
| pico540_pico540.012 | 0.675085314 | 143 | 0.67508531 | 157 | 22451 | generalists |
| pico540_pico540.027_sub | 0.723747451 | 149 | 0.72374745 | 167 | 24883 | generalists |
| pico540_pico540.18 | 0.651685597 | 136 | 0.6516856 | 149 | 20264 | generalists |
| pico540_pico540.27 | 0.817286762 | 178 | 0.81728676 | 190 | 33820 | generalists |
| pico540_pico540.28 | 0.865326377 | 187 | 0.80893206 | 187 | 34969 | generalists |
| pico540_pico540.38 | 0.739865506 | 155 | 0.72652938 | 168 | 26040 | generalists |
| pico540_pico540.5 | 0.501966447 | 92 | 0.42111139 | 68 | 6256 | potential_specialists |
| pico550_pico550.028 | 0.895886411 | 193 | 0.79539277 | 182 | 35126 | generalists |
| pico550_pico550.037 | 0.757818404 | 158 | 0.66745176 | 154 | 24332 | generalists |
| pico550_pico550.14 | 0.843786929 | 183 | 0.80327018 | 185 | 33855 | generalists |
| pico550_pico550.28 | 0.473403806 | 84 | 0.47340381 | 99 | 8316 | potential_specialists |
| pico550_pico550.45 | 0.788307111 | 167 | 0.77279984 | 175 | 29225 | generalists |
| pico550_pico550.49 | 0.386092589 | 35 | 0.38609259 | 53 | 1855 | specialists |
| pico551_pico551.013 | 0.884680099 | 192 | 0.78746679 | 179 | 34368 | generalists |
| pico551_pico551.11 | 0.804669956 | 174 | 0.70581874 | 165 | 28710 | generalists |
| pico551_pico551.14 | 0.736842105 | 153 | 0.52631579 | 113 | 17289 | potential_specialists |
| pico551_pico551.4 | 0.801339111 | 172 | 0.77648755 | 177 | 30444 | generalists |
| pico552_pico552.021 | 0.384400725 | 34 | 0.38440073 | 52 | 1768 | specialists |
| pico552_pico552.026 | 0.919191039 | 197 | 0.7931623 | 181 | 35657 | generalists |

